# Non-canonical *Staphylococcus aureus* pathogenicity island repression

**DOI:** 10.1101/2021.09.07.459249

**Authors:** Laura Miguel-Romero, Mohammed Alqasmi, Julio Bacarizo, Jason A. Tan, Richard J. Cogdell, John Chen, Olwyn Byron, Gail E Christie, Alberto Marina, José R Penadés

**Author notes:** Corresponding authors: José R Penadés, MRC Centre for Molecular Bacteriology and Infection Imperial College London,; Alberto Marina, Instituto de Biomedicina de Valencia (IBV-CSIC). These authors contributed equally to this work (Laura Miguel-Romero, Mohammed Alqasmi). Department of Biology, Augsburg University, Minneapolis, MN 55454 USA.

## Abstract

Mobile genetic elements (MGEs) control their life cycles by the expression of a master repressor, whose function must be disabled to allow the spread of these elements in nature. Here we describe an unprecedented repression-derepression mechanism involved in the transfer of the *Staphylococcus aureus* pathogenicity islands (SaPIs). Contrary to the classical phage and SaPI repressors, which are dimers, the SaPI1 repressor Stl^SaPI1^ presents a unique tetrameric conformation, never seen before. Importantly, not just one but two tetramers are required for SaPI1 repression, which increases the novelty of the system. To derepress SaPI1, the phage-encoded protein Sri binds to and induces a conformational change in the DNA binding domains of Stl^SaPI1^, preventing the binding of the repressor to its cognate Stl^SaPI1^ sites. Finally, our findings demonstrate that this system is not exclusive to SaPI1 but widespread in nature. Overall, our results characterise a novel repression-induction system involved in the transfer of MGE-encoded virulence factors in nature.

**Significance:** While most repressors controlling the transfer of mobile genetic elements are dimers, we demonstrate here that the Staphylococcal pathogenicity island 1 (SaPI1) is repressed by two tetramers, which have a novel structural fold in their body that has never been seen before in other proteins. Moreover, by solving the structure of the SaPI1 repressor in complex with its inducing protein Sri, we have demonstrated that Sri forces the SaPI1 repressor to adopt a conformation that is incompatible with DNA binding, explaining how SaPI1 is induced. Finally, our results demonstrate that this repression system is not exclusive of the SaPIs but widespread in nature. Our studies provide important insights understanding how SaPIs spread in nature.

## INTRODUCTION

*Staphylococcus aureus* pathogenicity islands (SaPIs) are the prototypical members of a widespread family of mobile genetic elements (MGEs), the phage-inducible chromosomal islands (PICIs; 1). SaPIs are clinically important because they encode and disseminate toxin and antibiotic resistance genes (1, 2). Normally, these elements reside passively integrated into the bacterial chromosome, thanks to the expression of a master repressor, called Stl (<u>S</u>aPI transcription leftward; 3). Unlike most phage repressors, the SaPI-encoded Stl repressors are not cleaved after induction of the cellular SOS response. This is because the Stl repressors are insensitive to the activated RecA protein. Instead, SaPI activation depends on the formation of a complex between Stl and specific phage proteins, which act as inducers for the SaPI cycle (4–6). Since SaPIs require the phage machinery for packaging (7, 8), this strategy ensures that SaPIs will be induced only in the presence of their prey, the phages.

Different SaPIs encode different repressors, and therefore require different phage proteins as inducers. Thus, the inducers for SaPIbov1, SaPI1 or SaPI2 are the phage-encoded dUTPases, Sri or recombinase proteins, respectively (4–6). Because of the singularity of the induction-repression mechanism present in the SaPIs, and to gain more insight into this interesting system, we recently solved the structure of the SaPIbov1 Stl (Stl^SaPIbov1^) repressor alone and in complex with two different inducing proteins, the dimeric and trimeric dUTPase proteins encoded by phages O11 and *ϕ*11, respectively (9). Our studies revealed that Stl^SaPIbov1^ is a canonical dimer, with a modular structural organisation reminiscent of that previously reported for many phage- and other MGE-encoded repressors, including the CI repressor from the archetypical phage *λ* (9, 10). These repressors have an N-terminal domain responsible for recognising and binding to their cognate DNA operator regions, and a C-terminal domain necessary for repressor dimerisation and inducer recognition. Since the dimer is absolutely required for repression, phages and other MGEs that are sensitive to the SOS response encode repressors that are structural homologs of LexA and therefore, also undergo self-cleavage when RecA is activated (SOS is induced). The repressor self-cleavage disrupts the dimer, activating either the lytic cycle of the prophages or the transfer of the different MGEs (11, 12). In the case of SaPIbov1, the only SaPI structurally characterised so far, the Stl^SaPIbov1^ dimer is also disrupted, a process that in this case occurs after the interaction of Stl^SaPIbov1^ with its cognate phage-encoded inducer proteins (9).

Our previous result suggested that the mechanism of action observed for Stl^SaPIbov1^ was similar to that observed for the classical CI-like repressors, except for how the dimers are disrupted in each scenario. In the case of SaPIbov1, this difference just represents the evolutionary adaption that this element has realised to sense the presence of its inducing phage by recruiting additional domains that play the dual role of mediating dimerisation and interacting with the inducer. However, the existence of multiple Stl repressor proteins raised the question whether this mechanism of action is conserved in all the SaPI repressors, or conversely, whether they sense their helper phages via as yet unidentified strategies. To answer this question, we analysed SaPI1. This is one of the prototypical islands used to decipher the biology of the SaPIs and is clinically relevant because it encodes the TSST-1 toxin, responsible for a rare but important human disease, toxic shock syndrome (13). Two additional factors reinforced the use of SaPI1 as a model. Firstly, Sri, its anti-repressor protein, is a small protein of 9 kDa, which raises the question of how this small protein de-represses SaPI1. Secondly, and contrary to what is seen with SaPIbov1 and other prophages, our unpublished results indicate that the repression mediated by the SaPI1 Stl (Stl^SaPI1^) is extremely strong. Thus, while some SaPIs – such as SaPIbov1 or SaPIbov2 – exhibit some level of basal excision from the bacterial chromosome in the absence of helper phage (14), SaPI1 remains integrated, suggesting that this island has a different repression system.

Here we confirm this idea and present a new repression-derepression system involved in the control and transfer of MGEs. Importantly, our results demonstrate that this new system is not exclusive to the SaPIs but widespread in nature. Once again, SaPIs provide an example of biological surprises, involved in promoting the emergence of novel bacterial virulent clones.

## RESULTS

### SaPI1 Stl is a tetramer

To understand the molecular basis of SaPI1 Stl (Stl^SaPI1^) mechanism of action we initiated these studies with the intention of solving its atomic structure, alone or bound to its cognate DNA. Although this was not possible, during its purification chromatographic assays revealed that Stl^SaPI1^ is a tetramer in solution. While the Stl^SaPI1^ monomer has 244 residues and a predicted molecular weight of 29 kDa (33 kDa after taking into account the N-terminal His (6) tag added to facilitate its purification), in size-exclusion chromatography (SEC) it eluted in a single peak as a protein of 133 kDa, which corresponds to four molecules of Stl^SaPI1^ (Fig. S1). This result was interesting, not just because most of the characterised MGE-encoded repressors are dimers, but also because it raised interesting questions about how Stl^SaPI1^ performs its function and how Sri, the SaPI1 inducer protein, promotes SaPI1 induction. Our initial hypothesis was that Sri would disrupt the Stl^SaPI1^ oligomeric state, as was observed for the Stl repressor of SaPIbov1 (Stl^SaPIbov1^) where dUTPases (both dimeric and monomeric) induce repressor monomerisation from its functional dimer. However, the SEC analysis of the Stl^SaPI1^-Sri complex ruled out this possibility and demonstrated that the tetrameric form is not affected by the presence of the inducing protein (Fig. S1).

This was confirmed by the X-ray structure of Stl^SaPI1^ in complex with the Sri protein from phage 80*α* (Fig. 1). The structure of the Sri-Stl^SaPI1^complex (Sri-Stl^SaPI1^) was determined to 2.9 Å resolution by single-wavelength anomalous dispersion (SAD) by using the diffraction of a selenomethionine-substituted derivative crystal (Table 1). The asymmetric unit of the crystal contained two Stl^SaPI1^ monomers (subunits A and B) in a dimeric organisation around a non-crystallographic two-fold axis with one Sri molecule binding to the Stl^SaPI1^ subunit A (Fig. 1B). The electron density observed in the model suggested that a second Sri molecule binds to the corresponding position of Stl^SaPI1^ subunit B. However, this additional Sri molecule showed an extremely low occupancy that allowed us to trace the main chain of fewer than 30% of its residues. In correlation with the low occupancy of the second Sri molecule, two regions of Stl^SaPI1^ subunit B (residues 33-47 and 85-96), which mediate interactions with the inducer in subunit A, present high flexibility that prevents its tracing. Similarly, we were unable to trace the N-terminal six residues, which are also placed in this area on subunit A, supporting the idea that the presence of Sri stabilises the Stl^SaPI1^ N-terminal region. In agreement with the size exclusion experiments, assembly analysis with PDBePISA server (15) indicates that the Stl^SaPI1^ dimer forms a stable tetramer (a dimer of dimers) exploiting the crystallographic two-fold axis. Therefore, the biological assembly of Sri-Stl^SaPI1^ is a box-shape heterooctamer of dimension 95 x 95 x 45 Å, and containing two Stl^SaPI1^ homodimers (subunits A-B and A*-B*) and four Sri molecules, two of them tightly bound (subunits A and A*) and two others with partial occupation (subunits B and B*).

**Figure 1.**
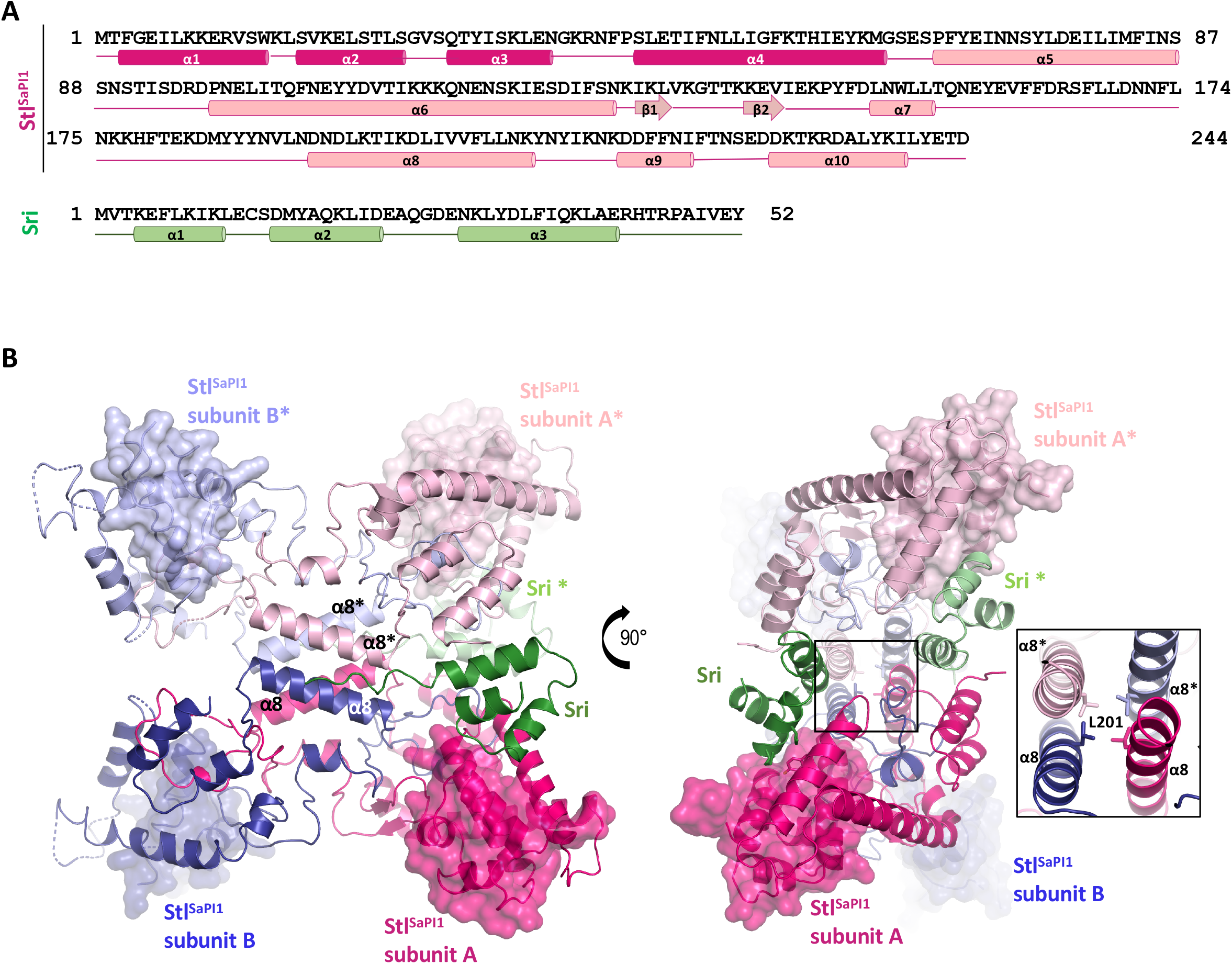
Structure of the Stl^SaPI1^–Sri complex. (A) Representation of the secondary structure of the Stl^SaPI1^ (in pink) and 80⍺ Sri (in green) proteins. The four first α helices of the Stl^SaPI1^ DBDs are colored in dark pink. (B) Two different views of the Stl^SaPI1^–80⍺ Sri complex structure. The molecules present in the symmetric unit of the crystal are coloured in dark pink and blue for Stl^SaPI1^ and green for Sri, while the asymmetric molecules are coloured in light colours. The Stl^SaPI1^ DBD surfaces are highlighted. The right part of the figure shows the Stl L201 residue located in the helix α8, its side-chain is represented as sticks.

**Table 1.** Data collection and refinement statistics.

| Table 1 Data collection and refinement statistics |  |
| --- | --- |
|  | Stl<sup>SaPI1</sup> - 80α Sri |
| **Data collection** |  |
| Space group | P6<sub>1</sub>22 |
| Cell dimensions *a*, *b*, *c* (Å) | 100.124, 100.124, 298.507 |
| α, β, γ (°) | 90, 90, 120 |
| Wavelength (Å) | 0.979 |
| Resolution (Å) | 75-2.9 |
| R<sub>merge</sub> (%) | 0.418 |
| Mean I/σ (I) | 15.3 |
| Completeness (%) | 98.6 |
| Redundancy | 5.2 |
| R<sub>pim</sub> | 0.038 |
| CC 1/2 | 0.998 |
| **Refinement** |  |
| Resolution (Å) | 75-2.9 |
| No. of reflections (observed/unique) | 20192/1019 |
| R<sub>work</sub>/R<sub>free</sub> (%) | 0.27/0.32 |
| Number of atoms | 4449 |
| Protein | 4284 |
| Water | 108 |
| Others | 55 |
| Bond lengths (Å) | 0.0053 |
| Bond angles (°) | 1.396 |
| **PDB ID code** | 7P4A |

Each Stl^SaPI1^ subunit presents a N-terminal DNA binding domain (DBD) with a conserved HTH XRE motif formed by 4*α* helices (helix *α*1-*α*4; residues 1-67) (Fig. 2A). This domain is quite similar to that found in the Stl^SaPIbov1^ structure (PDB ID 6H49) or in many phage repressors, including the phage *λ*CI (PDB ID 1LMB), showing the superimposition of Stl^SaPI1^ DBD with the equivalent C*α* atoms of these repressors with root-mean-square deviations (RMSD) of 1.40 and 2.3 Å, respectively (9, 16)(Fig. S2).

**Figure 2.**
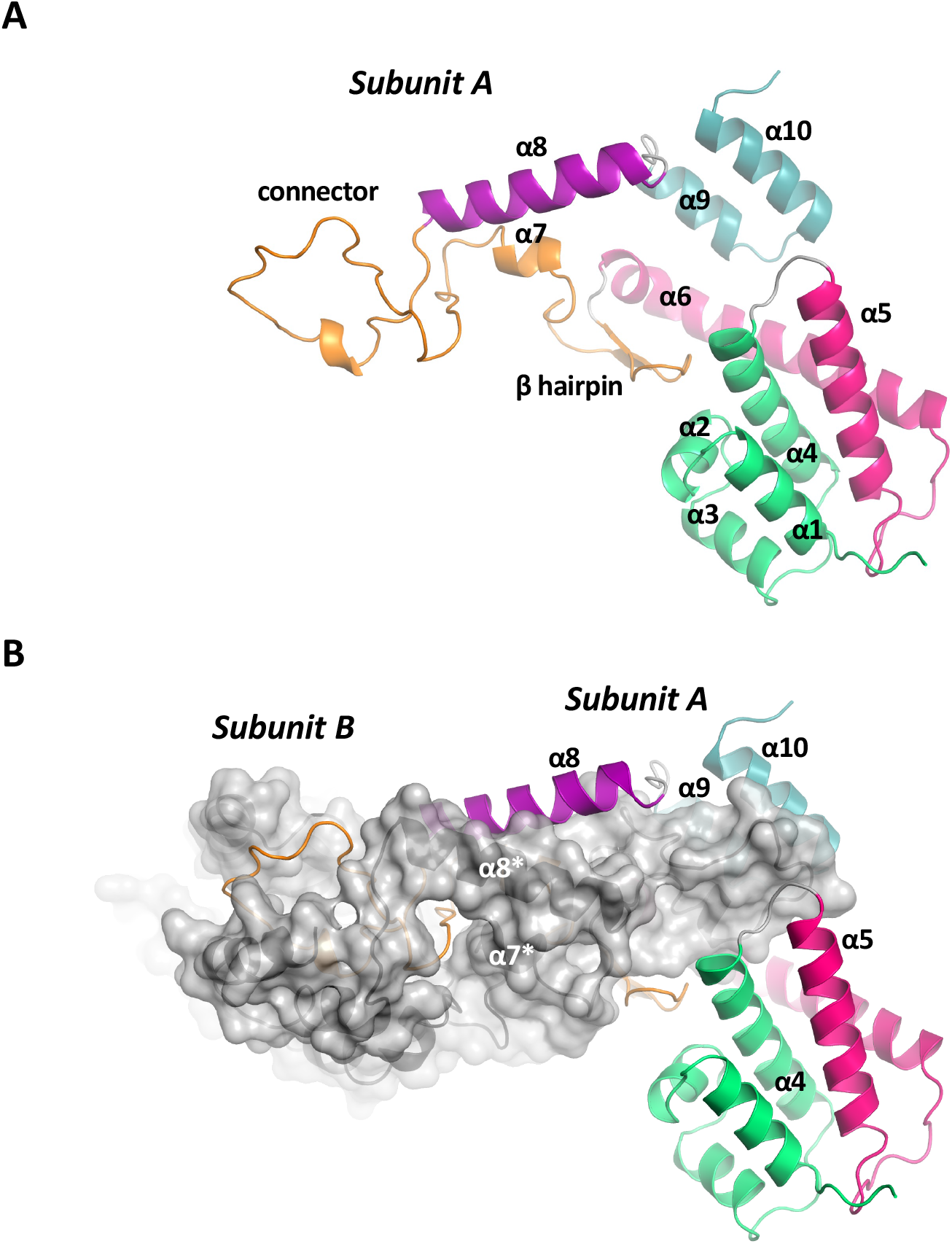
Structure of the Stl^SaPI1^ monomer and Stl^SaPI1^ dimer. (A) Structure of the Stl^SaPI1^ subunit A obtained from the Stl^SaPI1^-Sri complex. The DBD is coloured in green, the helices ⍺5 and ⍺6 are in pink, the central part of the molecule (β hairpin, ⍺7 and ⍺7- ⍺8 connection) is in orange. The helix ⍺8 is in purple and the C-terminal part (helices ⍺9 and ⍺10) is in blue. (B) Details of the Stl^SaPI1^ dimerization in the asymmetric unit of the crystal. One Stl^SaPI1^ molecule (subunit A) is coloured as defined in panel A while the second molecule of the dimer (subunit B) is grey and surface represented. Helices *α*4, *α*5, *α*9 and *α*10, plus the *β* harpin of subunit A create a cavity where the connector helices *α*7* and *α*8* from subunit B are located.

Conversely, no similar structures were found in the PBD database for the remaining body of the Stl^SaPI1^ repressor. The search of 3D homologs by comparison servers such as DALI or PDBefold (17, 18), using as a prey the structure formed by Stl^SaPI1^ residues 68-244, did not identify any structural homolog to Stl^SaPI1^. This unique architecture is composed of two long *α* helices (*α*5 and *α*6), with a long loop between them, which connect the DBDs to the central part of the protein. This central part is formed by two antiparallel strands (*β*1- *β*2) in a *β* hairpin and a short *α* helix (*α*7), followed by an extended region without secondary structure (residues 157-191) that projects from one subunit onto the other, returning to the same subunit via a long helix (*α*8). Finally, two shorter helices (*α*9 and *α*10) form the C-terminal part of the protein (Fig 2A). In this way, each subunit is highly elongated with a distance of more than 80 Å between the DBD domain and the region without secondary structure, which occupy both ends of the molecule and are connected mainly through helices *α*7 and *α*8.

In support of this novel structure, Stl^SaPI1^ self-associates in a conformation not previously observed in other repressors, with more than 40% of its residues interacting to form a huge dimerisation surface of ∼9230 Å^2^ per subunit (see Table S1 for a detailed description of the interaction between the two subunits). In the dimer, helices *α*4, *α*5, *α*9 and *α*10, plus the *β* harpin of one subunit creates a cavity where the connector helices *α*7* and *α*8* (residues 154-183) from the other subunit are located thereby embracing one monomer with the other (Fig. 2B). Although a huge number of contacts maintain the Stl^SaPI1^ dimer, the *α*7-*α*8 connector assembles the most important interactions supporting dimer formation (Table S1).

Analysing the tetrameric state of Stl^SaPI1^ in more detail, we observed that the tetramer presents a more reduced oligomerisation interface that buries only ∼2150 Å^2^ of the tetramer surface (∼540 Å^2^ per subunit). This tetramerisation surface is generated mainly by the mutual interaction of helices *α*8 (residues 193-208) from each subunit forming an antiparallel four-helix bundle in the tetramer (Fig. 1B). Interactions are mainly hydrophobic and provided by the side-chains of residues N193, D194, T197, L201, V204, F205, N208 and K209, which face the centre of the tetramer (Table S2).

### Stl^SaPI1^ tetramerisation is necessary for SaPI1 repression

To gain more insight into the biology of the Stl^SaPI1^ repressor, we analysed whether the tetramer was required for SaPI1 repression. Our previous structural analysis suggested the importance of residue L201 for Stl^SaPI1^ tetramerisation since their mutual interaction projecting from the middle of *α*8 nucleates the hydrophobic core (Fig. 1B). In support of this, a Stl^SaPI1^ repressor carrying the L201E mutation (Stl^SaPI1 L201E^), which introduces a charged residue into this hydrophobic environment, was a dimer in solution (Fig. S1), confirming the role of this helix in Stl^SaPI1^ tetramerization. Since this mutant was still able to bind to Sri (Fig. S1), this confirmed that the Stl^SaPI1^ L201E mutation affected tetramer formation but not the overall structure of the dimer.

Next, we generated a plasmid in which the *β*-lactamase reporter gene was fused to *xis*, downstream of *str* and the Stl-repressed *str* promoter, and which also encoded the divergently- expressed *stl* (see scheme in Fig. 3A). As a control, we generated a derivative plasmid expressing Stl^SaPI1 L201E^. These plasmids were introduced into strain RN4220 lysogenic for the SaPI1 helper phage 80*α* (5), and the prophage was induced using mitomycin C (MC). Samples were then taken at different time points, and the expression of the *β*-lactamase reporter quantified. In accordance with previous studies (5, 6), in the absence of prophage induction, Stl^SaPI1^ blocked the expression of the *β*-lactamase reporter. MC induction of the resident 80*α* prophage, leading to the concomitant expression of the Sri inducer, de-repressed the system promoting the expression of the reporter gene (Fig. 3B). In contrast, the plasmid expressing the dimeric Stl^SaPI1 L201E^ repressor showed an extremely high expression of the reporter gene, even in the absence of prophage induction (Fig. 3B), confirming its inability to repress the system. Note that we tried to generate a SaPI1 derivative island encoding the Stl^SaPI1^ L201E mutation but this was impossible, since mutations affecting Stl function are lethal for the bacteria as a consequence of the uncontrolled replication of the SaPI (3), confirming the incapacity of this mutant to repress SaPI1. In summary, the aforementioned results clearly demonstrate that the tetrameric structure is absolutely required for Stl^SaPI1^ repression.

**Figure 3.**
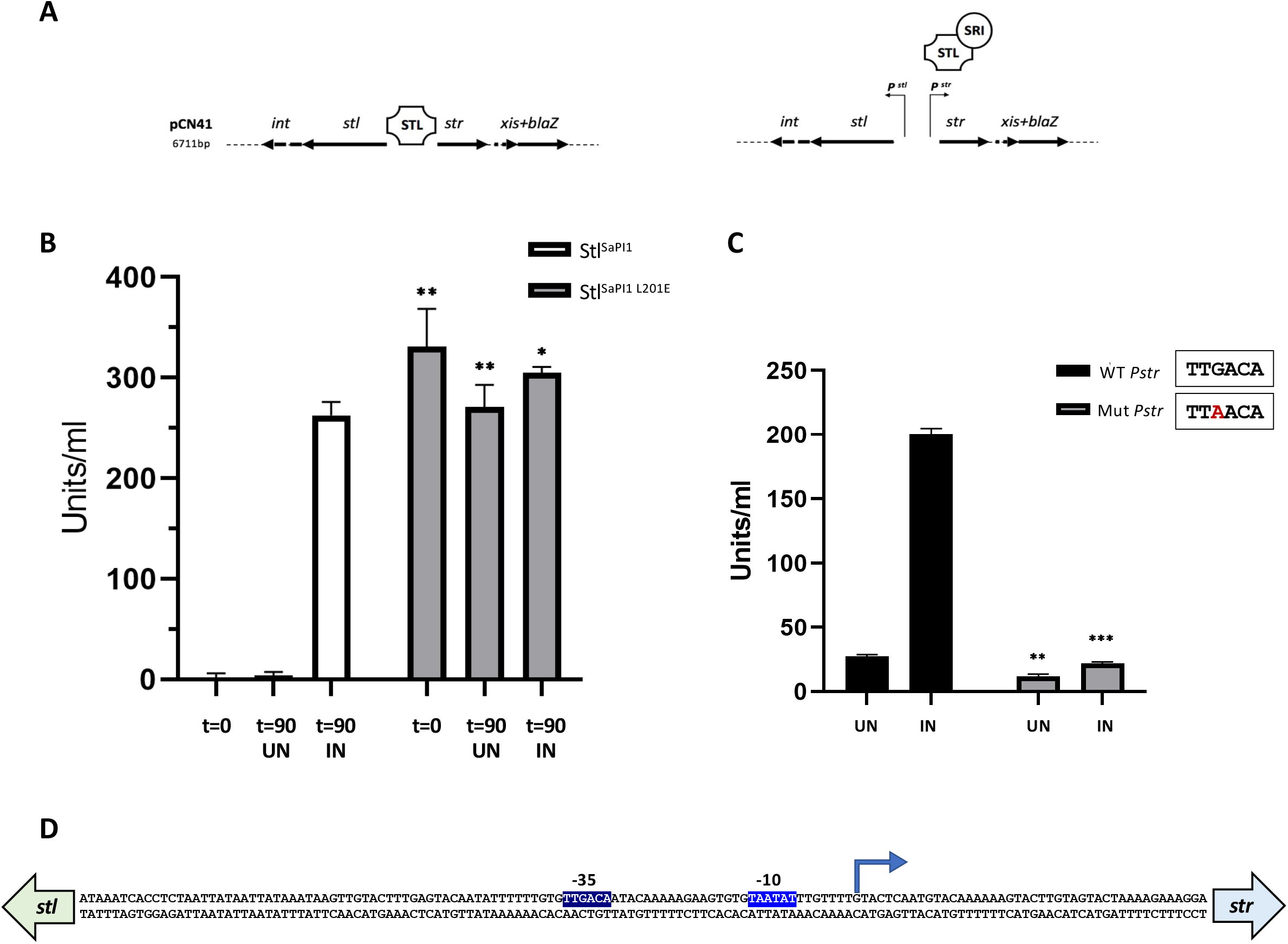
The Stl^SaPI1^ tetramer is required for SaPI1 repression. (A) Schematic representation of the pCN41 derivative reporter plasmid used to analyse Stl^SaPI1^ repression. In the absence of phage induction, expression of the *str*, *xis* and *blaZ* genes is repressed by Stl^SaPI1^. Induction of the 80α prophage results in expression of the SaPI1 inducer Sri, which binds to Stl^SaPI1^, promoting the expression of all the genes repressed by Stl^SaPI1^, including the *blaZ* reporter. (B) Lysogenic strains for phage 80α, carrying pCN41 derivative plasmids expressing either wt Stl^SaPI1^ or Stl^SaPI1^ L201E, were MC-induced (IN) or not (UN), and the expression of the *blaZ* reporter analysed at time zero (t=0) or 90 min (t=90). The means and standard deviations from three independent experiments are represented. A t-test comparation was performed to compare the different samples (WT vs mutant) taken at the same time points (*, P< 0.0332; **, P< 0.0021) (C) Characterization of the *str* promoter. Lysogenic strains for phage 80α, carrying pCN41 derivative plasmids containing either the intact intergenic region between *stl* and *str*, or a region carrying a point mutation in the putative -35 region of P*str*, were MC-induced (IN) or not (UN), and the expression of the *blaZ* reporter analysed as in (B) at 90 min. The means and standard deviations from three independent experiments are represented. A t-test comparation was performed to compare the the different samples (WT vs mutant) taken at the same time points (**, P< 0.0021; ***, P< 0.0002) (D) DNA sequence of the SaPI1 *stl-str* intergenic region. The P*str* +1 is represented with a blue arrow and the P*str* -10 and -35 sequences are highlighted in blue.

### Characterisation of the *stl* and *str* promoter regions

The next question we tried to answer is why the tetramer is required for SaPI1 repression. To do this, we initially performed 5’-RACE experiments to identify the *stl* and *str* promoter regions. While we were not able to identify the +1 for the *stl* transcript, we determined the start of *str* transcription which allowed us to identify the -10 and -35 regions present in the *str* promoter (Fig. 3D). This promoter showed the canonical sequence recognised by the RNA pol, suggesting that this is a strong promoter (19). To validate this result, we made use of the aforementioned *β*-lactamase reporter plasmid and introduced a single point mutation in the putative *str* -35 region (see scheme in Fig. 3C). The wild type (wt) and mutant plasmids were introduced into the strain lysogenic for phage 80*α*, and the expression of the *β*-lactamase reporter gene, which is fused to *xis*, was analysed after 90 min MC induction of the 80*α* prophage. The mutation introduced in the -35 region of the *Pstr* promoter completely abolished *str* expression, confirming that this region is essential for *Pstr* function (Fig. 3C).

### The SaPI1 *stl-str* intergenic region contains 8 Stl^SaPI1^ binding sites

The existence of 4 DBDs in the Stl^SaPI1^ tetramer, together with the fact that the dimeric Stl^SaPI1 L201E^ does not repress the island, raised two additional questions: how many boxes are present in the DNA region recognised by Stl^SaPI1^, and how are they organised? Foot-printing experiments using Stl^SaPI1^ and its cognate SaPI1 *stl-str* DNA intergenic region revealed the presence of two protected regions in both the top and bottom strands, separated by 24 bp (Fig. 4A). We used this region because in all characterised SaPIs the Stl proteins bind to the region that lies between *stl* and *str*(5)(9) and our reporter assays showed as regulated by Stl. The detailed analysis of the protected regions identified 8 putative Stl binding sites, organised as 4 different putative operators (1–4) (Fig. 4B). Operators 3 and 4 appear to represent a higher affinity site. This region is fully protected at a lower concentration of Stl than the region containing operators 1 and 2, and requires a higher concentration of Sri to lose protection. Each of these putative operators shows an almost perfect palindromic organisation containing two inverted 6 bp repeats with the consensus sequence TGTACT (called boxes A and B) separated by 3 bp (Fig. 4B). Importantly, operator 2 overlaps with the previously identified *Pstr* -35 region, suggesting that Stl^SaPI1^ represses expression of most of the SaPI1 genes by blocking the binding of the RNA polymerase to *Pstr*.

**Figure 4.**
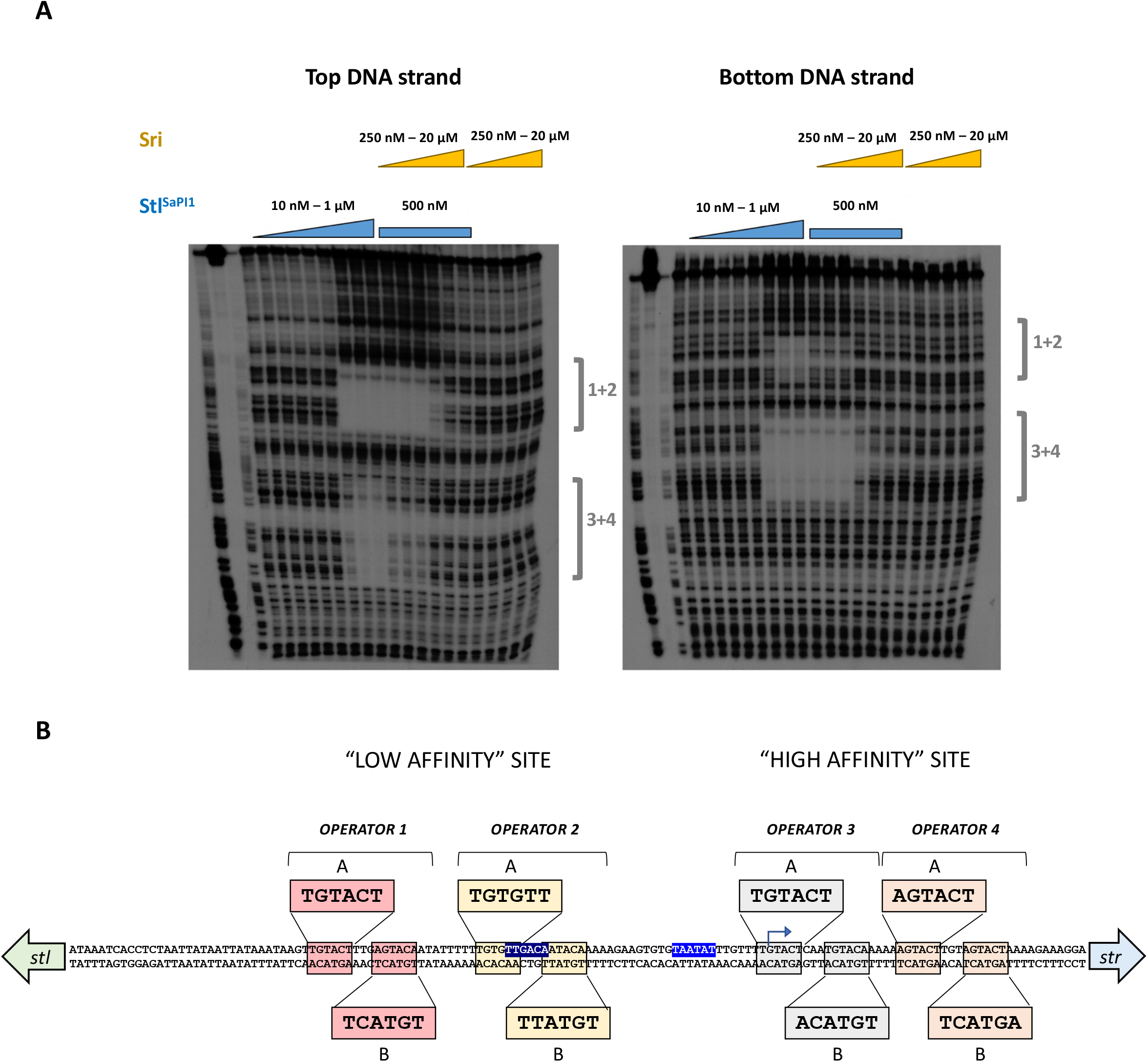
Identification of the Stl^SaPI1^ binding sites in the SaPI1 *stl-str* intergenic region. (A) Foot-printing experiments carried out with the SaPI1 *stl-str* intergenic region and the Stl^SaPI1^ protein, alone or in the presence of Sri. Protein concentrations used in the experiment are represented in the figure. The protective regions associated with the operators 1 and 2 (1+2) or 3 and 4 (3+4) are shown in the figure. Regions 1+2 and 3+4 correspond to regions with low and high affinity for Stl^SaPI1^, respectively. (B) Schematic representation of the 4 operator sites for Stl^SaPI1^. Both palindromic boxes of each operator are highlighted as A and B. The P*str* +1 is represented by a blue arrow. The P*str* -10 and -35 sequences are also highlighted in blue.

Since classical repressors containing HTH-XRE domains are usually dimers that bind to palindromic operators, the existence of 4 operators suggested that two Stl^SaPI1^ tetramers would bind to this region (corresponding to 4 classical dimers). To test this idea, we first confirmed that the 4 identified operators are functional, recognising and binding to the Stl^SaPI1^ DBDs. To simplify the interpretation of our results, we initially performed electrophoretic mobility shift assay (EMSA) experiments using the Stl^SaPI1 L201E^ mutant since the mutated residue is far away from the DNA recognition helix not affecting either the conformation of this domain or its ability to bind the operators. We hypothesised that each of the DBDs present in the Stl^SaPI1 L201E^ mutant would be able to bind as a dimer to each of the 4 operators independently. We analysed the binding of Stl^SaPI1 L201E^ to a set of DNA probes containing the 4 putative operators individually. As a control, one of the probes had all 4 putative operators mutated to A (Fig. 5C). As shown in Fig. 5A, when tested individually, Stl^SaPI1 L201E^ is able to recognise the 4 operators, showing highest affinity for operators 3 and 4, corroborating the foot-printing results.

**Figure 5.**
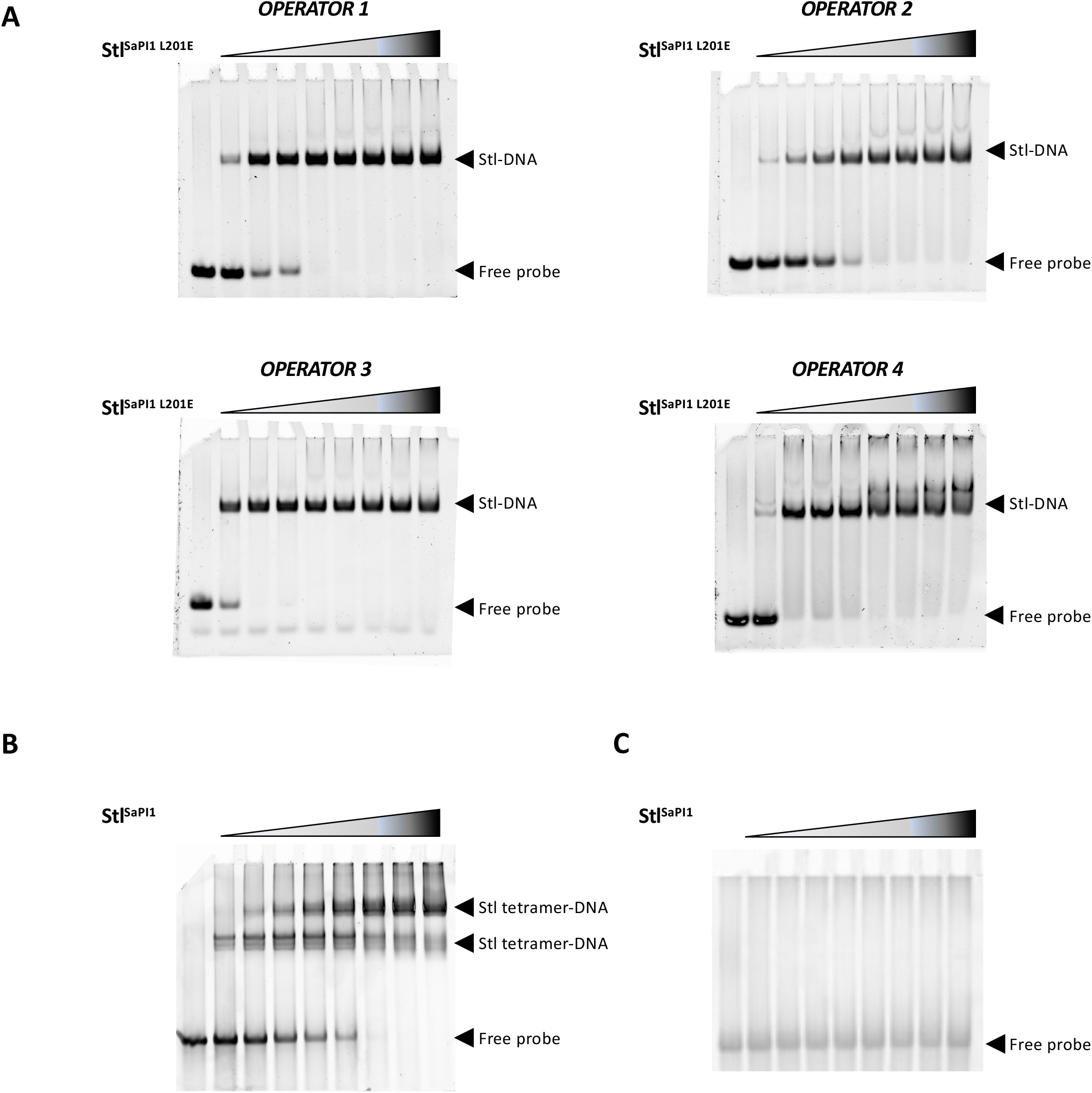
Characterization of the Stl^SaPI1^ binding sites present in the *stl-str* intergenic region. (A) The *stl-str* intergenic region contains 4 functional operator sites. EMSAs were performed using the Stl^SaPI1 L201E^ mutant protein (ranging from 0.5-4 µg per well), and DNA probes (1 µg) containing uniquely one putative operator site. (B) Two Stl^SaPI1^ tetramers bind to the *stl-str* intergenic region. The EMSA was performed using wt Stl^SaPI1^ (ranging from 1-8 µg per well) and a DNA probe (1 µg) containing the four operators present in the *stl-str* intergenic region. (C) As in (B), but is this case the four operator regions present in the DNA probe were mutated to AAAAAA.

### Two Stl tetramers bind to the *stl-str* intergenic region

Having demonstrated the existence of 4 operators in the *stl-str* intergenic region, we then analysed the interaction of the wt Stl^SaPI1^ with its cognate DNA region, using a DNA probe containing all 4 operators. In support of the hypothesis, our results demonstrated that two Stl^SaPI1^ tetramers bind to this DNA region, generating two different protein-DNA species, which could be easily differentiated by native gel electrophoresis (Fig. 5B).

Our previous results showed that Stl^SaPI1 L201E^ is unable to repress SaPI1 – even though it binds to the 4 Stl boxes. This result implied additional levels of complexity in the Stl^SaPI1^-DNA interaction. The existence of 4 operators with different affinities for Stl^SaPI1^, together with the requirement for 2 Stl^SaPI1^ tetramers, which show high flexibility for DBDs and the possibility of a large separation between DNA recognition *α*3 helices, prompted us to propose two different models for SaPI1 repression, which are currently under investigation (Fig. 6). In the first model, and as suggested by the foot-printing experiments, one tetramer would bind with high affinity to sites 3 and 4, while the other would bind to sites 1 and 2. In order to bind both sites at the same time, Stl must recognise an inverted palindrome of each operator that would generate a two canonical operators (3A-4B and 1A-2B) but in this case would be separated by 22-27 bp spacer instead of 3 bp (Fig. 6B). This long spacer would imply that the DNA-binding helices must be more than 60 Å apart to recognises the alternative operator, which should not be a problem for Stl^SaPI1^ since in the Sri-Stl^SaPI1^ structure shows that the *α*3 helices are separated by ∼70 Å in the dimer. Notice that although this type of separation between palindromes is unusual, it has recently been observed that the transcriptional activator AimR from *B. subtillis* phage SPbeta recognises an operator with similar organisation including a 25 bp spacer using highly flexible DBDs that are more than 75 Å apart (20). A protein-protein interaction between both tetramers could create a bigger protein-DNA complex able to stabilise Stl repression, which would explain why the Stl^SaPI1^ tetramer is required for SaPI1 repression. Another possibility could be that one dimeric part of the first tetramer would initially bind with high affinity to operator 3, and then the binding to operator 2 would induce a DNA torsion that is facilitated by the high T/A content of the inter-operator spacer. Next, a second tetramer would stabilize the protein-DNA complex by binding to operators 1 and 4 (Fig. 6B). This multiple interaction would be possible because of the high flexibility observed in the Stl^SaPI1^ DBDs (see below), and because of the ability of these domains to bind, bend and twist the DNA. We propose that either of these models stabilises the complex, making it impossible for the RNA polymerase to bind to the *stl* and *str* promoter regions. Although we have not yet confirmed either model, the binding of Stl^SaPI1 L201E^ to individual operators would support the latter model, while the footprinting data is more consistent with the former. Further studies will be needed to resolve this mechanism.

**Figure 6.**
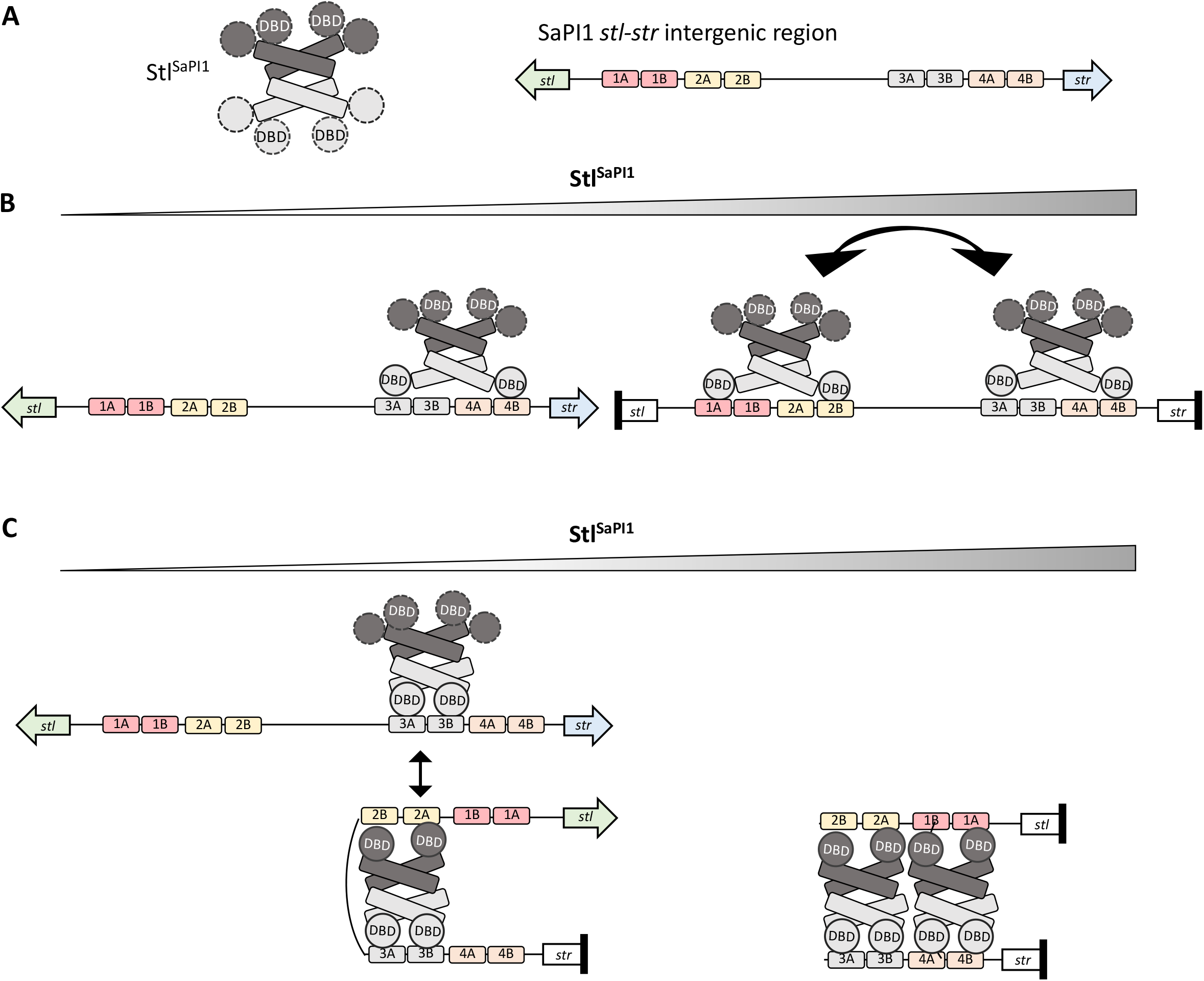
Proposed models for Stl^SaPI1^ repression. (A) Representation of the Stl^SaPI1^ tetramer and the SaPI1 *stl-str* intergenic region. The two dimers that from the Stl^SaPI1^ tetramer are dark and light grey and the mobility of the DBDs is represented by dashed lines. The 4 operators are represented with different colors and the inverted repeats are named as A and (B) Model 1. When Stl^SaPI1^ concentration is low, one tetramer binds the high affinity operators 3 and 4, which the DBDs from the dimer binding to the repetitions 3A and 4B. When Stl^SaPI1^ concentration increases, a second tetramer binds repetitions 1A and 2B form in a similar way than the tetramer binds to the operators 3 and 4. Since tetramer formation is required for SaPI1 repression, the two tetramers would interact to stabilize the complex. (C) Model 2. At low Stl^SaPI1^ concentration, one dimer of the Stl^SaPI1^ tetramer binds both repetitions (A and B) from operator 3, while the other binds to the repetitions present in operator 2, creating a torsion in the DNA, which favours *str* repression. When Stl^SaPI1^ concentration increases, a second tetramer in a similar manner operators 1 and 4, stabilizing the complex and increasing repression.

### Molecular basis of SaPI1 de-repression by Sri

Our results demonstrate that in each operator, two Stl^SaPI1^ DBDs bind to two palindromic sequences which are separated by 3 bp. However, in the X-ray crystallography structure of the SaPI^SaPI1^-Sri complex, the distance observed between the two DBDs (> 60 Å) is higher than that required to bind to the Stl boxes in the operators (30 Å, Fig. S3). Sri is a 52 residue, 9 kDa protein composed of 3 short *α* helices (*α*1-*α*3) and a 7 residues C-terminal tail. DALI search showed that Sri is structurally similar (1.28 Å RMSD over 41 C*α* superimposed) to the phage 77 ORF104, whose structure was previously solved in complex with DnaI, its cellular partner (21) (PDB ID 5HE9). When complexed with Stl^SaPI1^, Sri is inserted in the Stl^SaPI1^ tetramer, interacting with 3 of the 4 Stl^SaPI1^ subunits and burying ∼1550 Å^2^ of its surface, which corresponds to ∼33% of the Sri molecular surface. Each Sri molecule mainly interacts with the Stl^SaPI1^ DBD of one subunit by using helices *α*2 and *α*3 that contact with Stl^SaPI1^ helices *α*1, *α*4 and *α*5. In addition, the Sri *α*3 helix interacts with the C-terminal *α*10 helix of the symmetrically related subunit (A*) in the Stl^SaPI1^ tetramer and its extended C-terminal tail is positioned over the 2 *α*8 helices (subunits A* and B), which nucleate the Stl^SaPI1^ tetramerisation. By performing this network of interactions Sri glues the Stl^SaPI1^ DBDs to the main body of the Stl^SaPI1^ tetramer, restricting its conformational freedom. The fact that a low occupancy of Sri in the second binding site of the Stl^SaPI1^ dimer correlates with a high flexibility of the DBD supports this Sri function. Therefore, we hypothesise that Sri de-represses SaPI1 by fixing the Stl^SaPI1^ DBDs (Fig. 1, Table S3) in a conformation that is not compatible with DNA binding (Fig. S3).

To test this hypothesis, we first validated our structural data for the Stl^SaPI1^-Sri interaction. To do this, we mutated the Stl^SaPI1^ residue Y76 to A (Stl^SaPI1_Y76A^), which we anticipated was important for the Stl^SaPI1^-Sri complex stabilisation by projecting its side chain into a hydrophobic pocket generated by Stl^SaPI1^ residues W14, M63, F69, I72 and Y76 (Fig. 7A, Table S3). Pull-down assays confirmed that the Stl^SaPI1 Y76A^ repressor was unable to bind to Sri (Fig. 7B). To show that this mutation affected only the interaction with Sri, but not the ability of Stl^SaPI1 Y76^ to repress the island, we used the aforementioned *β*-lactamase reporter plasmid (see scheme in Fig 3A) expressing either the wt or the Stl^SaPI1 Y76A^ mutant repressor. These plasmids were introduced into the 80α lysogen and the expression from the Stl-repressed *str* promoter was measured after induction of the 80α prophage. Compared with that observed for the wt Stl^SaPI1^, no significant activity was observed in the plasmid expressing Stl^SaPI1 Y76A^ under all of the conditions tested, confirming that this mutant protein is still able to repress the island but is insensitive to the helper phage Sri protein (Fig. 7C). Finally, we tested the impact of the Stl^SaPI1Y76A^ mutation *in vivo*. We generated a SaPI1 *tst*::*tet*M derivative island expressing Stl^SaPI1 Y76A^.

**Figure 7.**
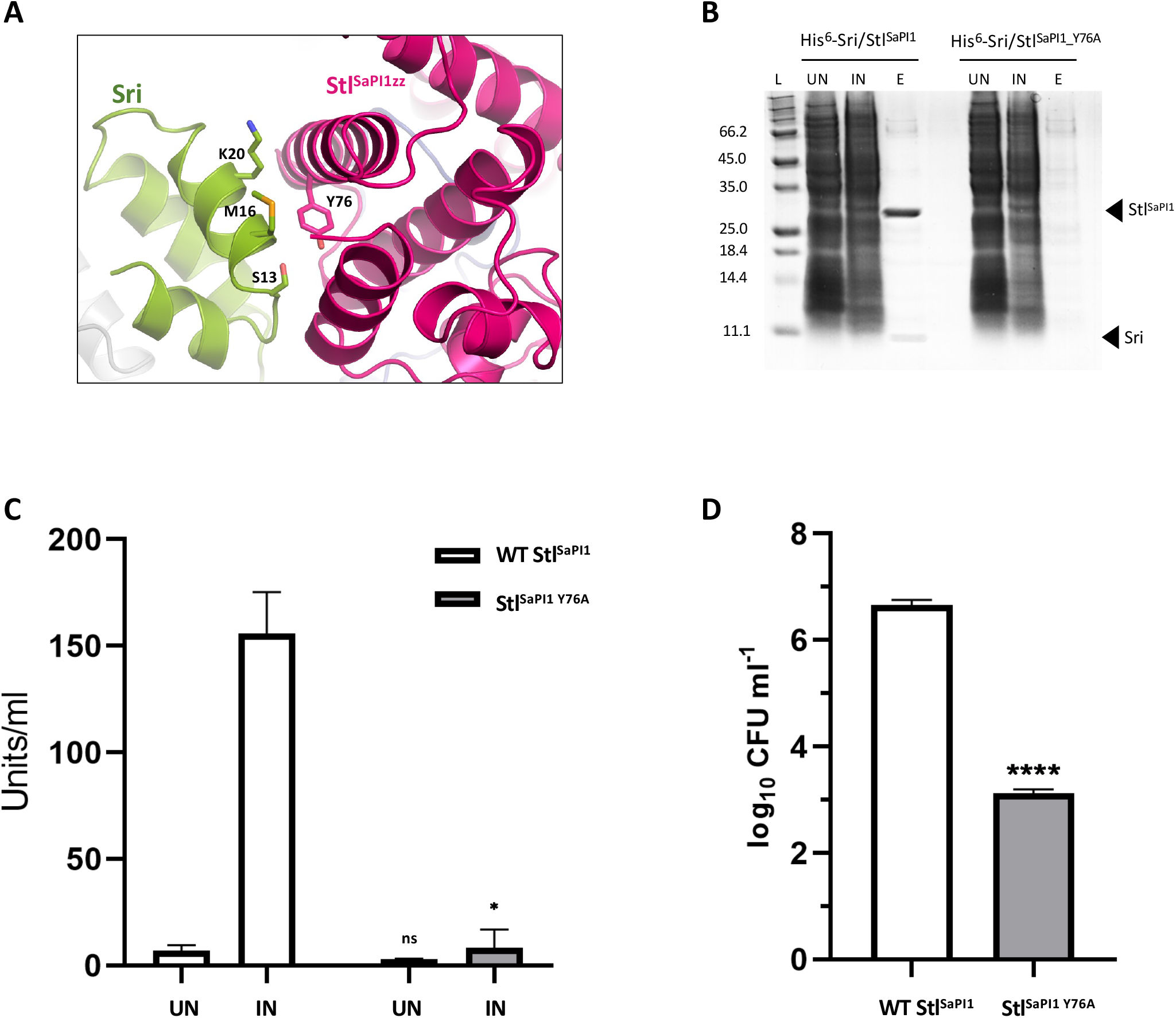
Characterization of the Stl^SaPI1^-Sri interaction. (A) Details of the Stl^SaPI1^ Sri interaction. The Stl^SaPI1^ Y76, and the Sri S13, M16 and K20 residue side-chains are represented as sticks. (B) SDS-PAGE gel after pull-down experiments in which the His(6)-tagged Sri protein was co-expressed either with the wt Stl^SaPI1^ or the Stl^SaPI1 Y76A^ mutant proteins. Uninduced (UN), induced (IN) and eluted (E) from the Ni^2+^ column. (C) Lysogenic strains for phage 80α, carrying pCN41 derivative plasmids expressing either wt Stl^SaPI1^ or Stl^SaPI1 Y76A^ were MC-induced (IN) or not induced (UN), and the expression of the *blaZ* reporter analyzed at 90 min. The means and standard deviations from three independent experiments are represented. A t-test comparation was performed to compare Stl^SaPI1 Y76A^ versus Stl^SaPI1^ (*, P< 0.0332; ns, P< 0.1234). (D) Lysogenic strains for phage 80α, carrying wt SaPI1 *tst*::*tet*M or a derivative SaPI1 *tst*::*tet*M carrying the Stl^SaPI1^ Y76A mutation, were MC- induced (IN) or not induced (UN), and the transfer of the island quantified. The means and standard deviations from three independent experiments are represented. A t-test comparation was performed to compare Stl^SaPI1 Y76A^ versus (****, P< 0.0001).

**Figure 8.**
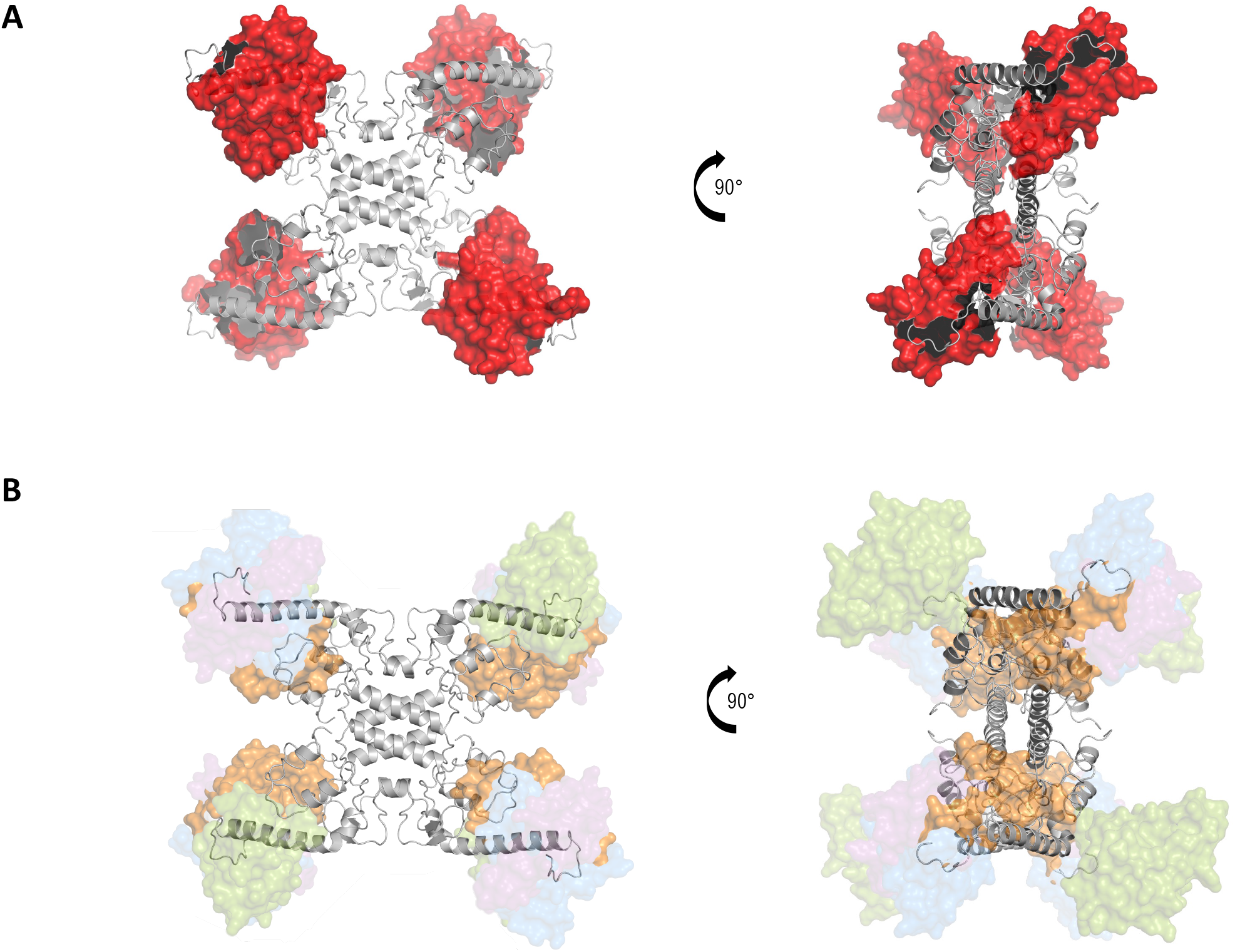
The Stl^SaPI1^ DBDs are flexible in solution. (A) AStl^SaPI1^ tetramer structure was generated by removing the Sri coordinates from the X-ray crystallography structure of the Stl^SaPI1^-Sri complex. Residues 90-244 are in grey and DBDs surfaces are in red. (B) Model with the DBD conformers which best fit to the SAXS data via the ensemble optimization method for Stl^SaPI1^ in solution. Stl^SaPI1^ residues 90-244 are in grey and the DBD conformers are represented in surface with different transparencies proportional to the percentage of that model in the total ensemble. The fit was obtained with ∼ 50% of model 1 (DBDs in orange), 25% of model 2 (DBDs in green), and 13% of models 3 (DBDs in blue) and 4 (DBDs in purple).

Note that this island carries an antibiotic resistance marker, which facilitates the transfer studies. The strain was then lysogenised with phage 80α, and the transfer of the island analysed after induction of the resident prophage. As a control we included the lysogenic strain for 80α carrying the wt SaPI1 *tst*::*tet*M. The transfer of the SaPI1 mutant island was significantly reduced compared with that observed for the wt SaPI1 (Fig. 7D). Taken together, these results validate the Stl^SaPI1^-Sri interactions revealed by the X-ray crystallographic data.

Next, although we were unable to obtain the Stl^SaPI1^ structure alone or in complex with its cognate DNA, we obtained low resolution structural information about this protein in solution using small-angle X-ray scattering (SAXS). First, we generated a SAXS data set of the same Stl^SaPI1^-Sri complex and for Stl^SaPI1^ alone merging 14 and 49 frames of SAXS data for which a constant Rg of 39.7 Å and 41.1 Å was estimated, respectively. The maximum particle dimension Dmax was 127.3 Å and 139.1 Å for Stl^SaPI1^-Sri complex and for Stl^SaPI1^ alone respectively (Fig. S4). The mass of Stl^SaPI1^-Sri monomer (including a 6-His tag on Sri) is 38997 Da and 29398 Da for Stl^SaPI1^ so the SAXS analysis was therefore strongly suggestive of a tetramer either of Stl^SaPI1^-Sri or Stl^SaPI1^ alone (155988 or 117592 Da respectively). The analysis of a model for a Stl^SaPI1^ tetramer, generated by removing the coordinates of Sri from the X-ray crystallography structure of SaPI^SaPI1^-Sri, indicates that the Stl^SaPI1^ tetramer in solution is more compact in complex with Sri than alone. Thus, the mobility of the Stl^SaPI1^ DNA-binding domains (DBDs) was modelled using EOM (22). 10000 models were generated in which the C-terminal domains (CTD, residues 101-247) were kept in the conformation observed by X-ray crystallography but the DBDs (residues 1-89) allowed to adopt positions consistent with their connection to the CTD via a native-like flexible linker (residues 90-100). In support of this, our crystallographic structure of Stl^SaPI1^-Sri showed high mobility in the Stl DBDs when Sri was weakly bound (subunits B and B*, Fig. 1B), suggesting that the structure of these DBDs is highly plastic in the absence of Sri, and therefore, compatible with binding to the DNA Stl boxes.

These results indicate that SaPI1 de-repression involves a mechanism different from that previously described for SaPIbov1 induction. While the latter involves separation of the Stl^SaPIbov1^ dimer by the inducing dUTPases (9), our biochemical, structural and *in vivo* data show that Sri de-represses SaPI1 by inducing a conformational change in the Stl^SaPI1^ DBDs, preventing the binding of these domains to their cognate Stl^SaPI1^ sites.

### Stl^SaPI1^ homologs are widespread in nature

Because of the unusual nature of the SaPI1 repression system, we wanted to know if it was exclusive to this island or more widespread in nature. In a search for Stl^SaPI1^-like homologues in the publicly accessible databases, different Stl^SaPI1^-like homologs were found, not just in other Staphylococci, but also in different species of *Bacillus* and *Virgibacillus*, which are 25- 35% similar to Stl^SaPI1^ (Table S4). Importantly, while the Stl^SaPI1^-like homologs present in the different *Staphylococcus* spp. were encoded by different members of the PICI family (Table S4), suggesting a mechanism of induction in common with that reported here for SaPI1, the homologs present in the other genera were encoded by MGEs other than PICIs, raising the question of how these elements are de-repressed. Moreover, the 3D models obtained for these proteins confirmed their high structural homology with Stl^SaPI1^ (TM-scores 0.6-0.7; Fig. 9A), and the detailed analysis of the secondary structure of these homolog repressors showed that the DBD (present in the first 3 *α* helices) and dimerisation (region between helices *α*7 and *α*8) or tetramerization (helix *α*8) key residues were extremely well conserved among these proteins (Fig. 9B). These results confirm the discovery of a new family of repressors involved in gene transfer and bacterial evolution.

**Figure 9.**
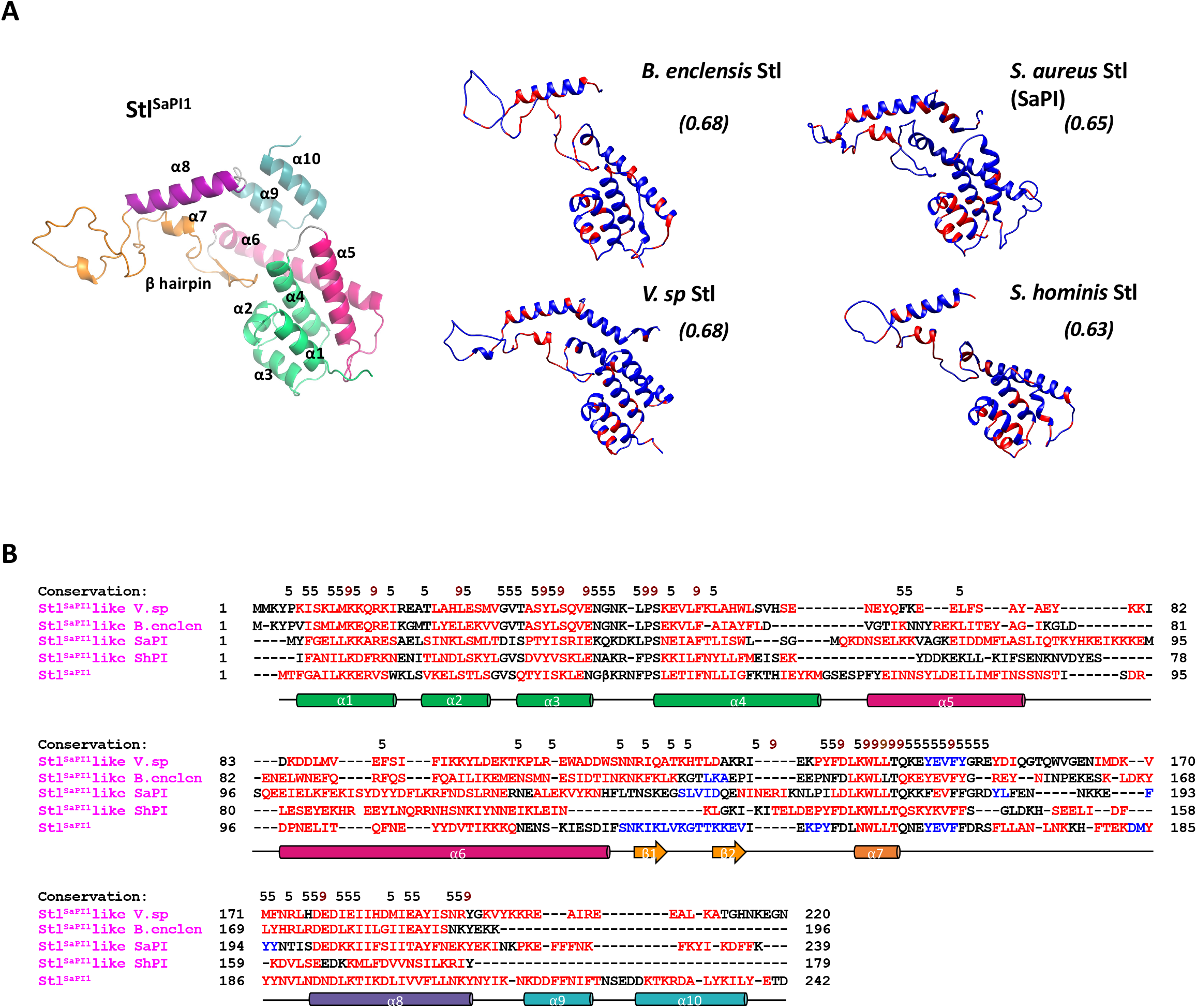
Stl^SaPI1^ repressor conservation in other species. (A) The Stl^SaPI1^ structure was used to model the other Stl^SaPI1^–like repressors present in other bacterial species. The colors in Stl^SaPI1^ are defined in Fig. 2A. The Stl^SaPI1^ residues conserved in the Stl^SaPI1-like^ repressors are colored in red, while the non-conserved residues are colored in blue. The TM-scores for the models are indicated in italics in brackets. (B) Structural alignment of Stl^SaPI1^ and Stl^SaPI1^–like repressors. The residues forming the ⍺ helices or the β strands are colored in red and blue, respectively. Residue conservation, 9 being conserved and 5 similar, is on the alignment. The Stl^SaPI1^ secondary structure is represented below the alignment. The colors of Stl^SaPI1^ secondary structure are defined in Fig 2A.

## DISCUSSION

Many MGEs, including prophages, PICIs or ICEs, control their life cycles by expressing master repressors that maintain these elements integrated into the bacterial chromosome. Importantly, the expression of these repressors must be precisely controlled, since an excess of the repressor would impede the induction and transfer of the element, while a reduced expression would generate either the loss or the activation of the element under unfavourable conditions.

To ensure this tightly regulated control, SaPI1 has evolved a unique system involving 2 Stl^SaPI1^ tetramers and 4 operators. As previously mentioned, we have proposed here two different models for SaPI1 repression (Fig. 6). The best characterised repressor so far is CI, from phage *λ*. CI represses both *cI* and *cro* expression by binding at operators *OL* and *OR*, each one composed of three repressor binding sites, named *OL1*, *OL2* and *OL3*, or *OR1*, *OR2* and *OR3*, respectively (23). Two CI dimers bind tightly and cooperatively to *OL1* and *OL2*, creating a tetramer that represses expression from the *pL* promoter (24). A similar tetramer bound structure is formed after the binding of two CI dimers to *OR1* and *OR2*, repressing in this case the expression from the *pR* promoter (24). To generate a more stable repression system, these two tetramers interact to form an octamer looping the DNA between the *OL* and *OR* operator regions (11, 25). While Stl^SaPI1^ and CI repression involve the formation of tetramers, these two systems are completely different, both structurally and mechanistically, probably in response to the different ways that phages and SaPIs are induced. In *λ*, the two tetramers and the octamer appear only after the binding of the CI dimers to their cognate binding sites, while Stl^SaPI1^ is always a tetramer. Moreover, our results indicate that the binding of the two Stl^SaPI1^ tetramers to their cognate DNAs is not cooperative but sequential. Structurally, these two repressors are completely unrelated, except in their DBD regions. Functionally, these two repressors also work in completely different ways. Thus, after activation of the bacterial SOS response, the RecA* protein will promote the autocleavage of CI, disrupting dimer, tetramer and octamer formation, while in the case of SaPI1, the Sri protein does not affect tetramerisation of Stl^SaPI1^ but will force the Stl^SaPI1^ DBDs to adopt a conformation that prevents their interaction with their cognate DNA boxes.

Another two systems involving tetramer formation have been described in the control of the transfer of different MGEs. In the first one, controlling the life cycle of the temperate *Salmonella* phage SPC32H (26), the repressor – here called Rep – can reversibly assemble into two oligomeric states, dimer and tetramer, in a concentration dependent manner. As with Stl^SaPI1^, Rep binds to the DNA as a tetramer, although these tetramers are structurally completely different. In fact, and contrary to what was seen with the Stl^SaPI1 L201E^ dimer, the dimeric Rep protein binds DNA weakly even at high concentrations. This difference can be easily explained since it has been proposed that in Rep the dimer pairs required for binding to the palindromic DNA sites originate from different dimers in the tetrameric Rep structure (26), while in the Stl^SaPI1^ they come from the same dimer. Another difference between these systems relates to how they are de-repressed. While the SaPI1 de-repressor Sri is a monomer that forces a conformational change in the Stl^SaPI1^ DBDs, the SPC32H anti-repressor Ant is a tetramer that binds to two dimeric Reps, breaking tetramer formation. Another important difference between the two systems is that only one tetramer is required for phage SPC32H repression, but two are required for SaPI1 repression (26).

The second system is found in the *Enterococcus faecalis* conjugative plasmid pCF10. This plasmid encodes PrgX, a repressor that blocks the expression of the genes involved in the conjugative transfer of this plasmid. As with Stl^SaPI1^, PrgX is a tetramer that binds to two different operator regions forcing a looping of the DNA (27). However, Rep, PrgX and Stl^SaPI1^ are completely unrelated structurally. Moreover, the mechanism involving pCF10 transfer is also different from that observed for SaPI1. The transfer of the pCF10 plasmid occurs in response to an intracellular pheromone signal, a peptide called cCF10 with the sequence LVTLVFV. As with Rep, binding of the cCF10 inducer to PrgX destabilises the PgrX tetramer, promoting conjugation (27).

The new repression mechanism described here for SaPI1 is possible due to the localisation of the Stl^SaPI1^ DBDs in the tetramer. Canonical members of the HTH-XRE family of repressors dimerise through their *α*5 helix. In Stl^SaPI1^, however, the *α*5 helix connects the DBDs with the rest of the protein by a long loop which confers upon them high mobility for DNA recognition and binding. The flexibility of the Stl^SaPI1^ DBDs was obvious when we compared the four DBD domains in our Stl^SaPI1^-Sri X-ray crystallographic structure versus the SAXS data obtained for the Stl^SaPI1^ protein alone. By interacting with Stl^SaPI1^ DBD helices *α*1, *α*4 and *α*5 and two other subunits in the tetramer, Sri maintains the Stl^SaPI1^ DBDs fixed in a conformation that prevents their binding to the operator regions present in the DNA. However, in the absence of Sri, the DBDs showed a more extended localisation that would allow them to interact with their cognate DNA boxes. This plasticity in the DBDs is likely to be the origin of our incapability to obtain X-ray structure crystallographic structures for Stl^SaPI1^ and Stl^SaPI1 L201E^ alone.

Another interesting feature of the new regulatory system described here relates to the fact that the primary role of the phage-encoded SaPI1 inducers is not to induce this island but to interact with the cellular DnaI protein (21), slowing down bacterial replication and facilitating phage reproduction. Thus, SaPI1 has evolved a repressor that uses a conserved phage protein as inducer. However, and since DnaI and Stl^SaPI1^ are completely unrelated both in sequence and structurally, how this small protein is able to interact with two unrelated proteins performing two different functions increases the interest of this system and is currently under study. Other interesting unsolved questions include what is the origin for this repressor, and how SaPI1 co-opted and evolved it to adapt to its life cycle requirement. While we cannot answer these questions yet, it is clear that this repressor is not unique to SaPI1 but present in other MGEs from different species. It is also clear the SaPIs have evolved an impressive arsenal of strategies to hijack the life cycle of their helper phages, making these elements one of the most sophisticated subcellular parasites in nature.

## MATERIAL AND METHODS

### Bacterial strains and growth conditions

Bacterial strains used in this study are listed in Table S5. Strains were grown at 37°C in Luria-Bertani (LB) agar or in LB broth with shaking (180 r.p.m.) for *E. coli* or in Tryptic Soy agar (TSA) or Tryptic Soy Agar broth (TSB) for *S. aureus*. Ampicillin (100 mg ml^-1^) or Tetracycline (20 mg ml^-1^; all Sigma-Aldrich) antibiotics were added when appropriate.

### Plasmid construction

The plasmids used in this study (Table S6) were constructed by cloning PCR products, amplified with the oligonucleotides listed in Table S7 (Sigma-Aldrich), into the appropriate vectors. The cloned plasmids were verified by Sanger sequencing (Eurofins Genomics).

### Protein Expression and Purification

Proteins were overexpressed from *Escherichia coli* BL21 (DE3) (Novagen) cells transformed with the corresponding expression plasmids (Table S6). Cultures were grown to an OD_600_ of 0.5–0.6 and protein expression was induced with 1 mM IPTG at 20°C for 16 h. Cells were harvested by centrifugation at 4°C, 4000 rpm for 20 min, resuspended in lysis buffer (100 mM Tris pH 8, 300 mM NaCl, 5ml Mg2Cl, 1mM ß-mercaptoethanol and protease inhibitor tablets (complete tablets, Roche) and lysed by sonication. The soluble fractions were obtained by centrifugation at 4°C, 15000 rpm for 30min and loaded onto a pre-equilibrated Nickel affinity column (Histrap 5ml; GeHealthcare). After wash with 20 mM, the proteins were eluted with lysis buffer containing 300 mM imidazole. The fractions containing the eluted protein were loaded in Superdex S200 26/600 and analyzed by SDS-PAGE and those fractions showing purest protein were selected, concentrated until 10-15mg/ml, and stored at -80°C. For Sri- Stl^SaPI1^ crystallization, the complex was digested with the TEV protease to eliminate the 6xHis tag at 4°C o/n in 100mM Tris-HCl pH8, 300mM NaCl and 5mM Mg2Cl buffer before loading it in gel filtration. Sri-Stl^SaPI1^ selenomethionine-labeled (SeMet) derivative complex was obtained using SelenoMethionine Medium Complete (Molecular Dimensions Ltd; MD 12-500), according to the manufacturer instructions, and purified as described previously.

### End-labeling SaPI1 *stl-str* DNA for footprinting experiments

SaPI1 *stl-str* DNA was amplified using *PfuTurbo* DNA polymerase (Agilent), dNTPs (Invitrogen), and primers GC83016C (5’- GTTCTTTAACCGAAAGCTTCCAACTCACTCTTTC- 3’) and GC83016B (5’-CAGTACGTTCGTGATACAAGCCATGTATTGATGTTC-3’). The PCR product was diluted 1:1,000 (approximately 0.001pmol/ µl) for preparing radiolabeled SaPI1 *stl-str* DNA. 5pmol of primer (5µl of 5 µM stocks) were end-labeled with 10 units of USB Optikinase (Affymetrix), and 7 µl of 6,000Cu/mmol ATP-ɣ32P (Perkin-Elmer) in 25 µl total volume. Incorporation of ATP-ɣ32P into primer was determined by TCA precipitation of a 5 µl aliquot of a 1:200 dilution and counting the precipitated material on filter paper using a scintillation counter. To obtain final radiolabeled SaPI1 *stl-str* DNA (*) used in footprint experiments, the remaining 24.5 µl of the undiluted Optikinase reaction (containing approximately 1pmol/ 5µl of either top or bottom-strand labelled primer) was added to the original 1:1,000 dilution of the SaPI1 PCR product (template), 1 µl of 10mM dNTP (Invitrogen), and 1 µl of *PfuTurbo* DNA polymerase (Agilent) for amplification in a final volume of 50 µl. Unincorporated primers were removed using QIAquick PCR Purification Kit (Qiagen) and eluted twice with 30 µl of water. To assess the final concentration of SaPI1 *stl-str* DNA*, a 1:200 dilution of the PCR cleanup material was made and dpm assessed using the same TCA precipitation/filter method and scintillation counter.

### G/A Ladder of SaPI1 *stl-str* DNA

The G/A ladder was generated using the piperidine method. First, 12µl of SaPI1 *stl-str* DNA* was added to 2µl of 1 mg/ ml salmon sperm DNA and 8 µl of Tris-EDTA (TE), pH 8 and incubated on ice. Second, 2µl of 4% formic acid made fresh from stock was added and incubated at 37°C for 45 minutes, and subsequently place back on ice. Third, 300 µl of Piperidine (Sigma-Aldrich) was added and incubated at 90°C for 30 minutes, and subsequently placed back on ice. Fourth, 10 µl of 5mg/ml salmon sperm DNA (50mg total) was added followed by gentle mixing. Fifth, 1ml of butanol was added, mixed, and centrifuged at approximately 20,000 rcf (x g) for five minutes; the top layer was carefully removed. A further 1.2ml of butanol was added, mixed, and centrifuged resulting in a pellet that contained precipitated DNA. Supernatant was removed and the pellet was gently washed with 150 µl of 1% SDS, followed by butanol precipitation (one 1ml, and two 0.5ml butanol precipitations). The DNA pellet was dehydrated with a speed vacuum, and resuspended in 20 µl of footprint loading buffer (0.5mL deionized formamide; 20µl of 0.25M EDTA, pH 7; 5µl XCFF and 5µl of BPB for visualization of sample migration during electrophoresis) and stored at -20°C. Lastly, 1:7.5 and 1:15 dilutions of the G/A ladder were made using the same footprint loading buffer in order to obtain exposure intensities more suitable for sequence determination of the Stl footprints on SaPI1 *stl-str* DNA.

### DNase I Footprinting

Footprinting reactions were carried out in EMSA buffer (PBS, pH 7.3, 75mM NaCl, 5mM MgCl2,1mM DTT, 0.1mg/ml BSA, 5% glycerol) supplemented with 2mM CaCl2, which is required for DnaseI activity, and 100ng/ µl poly(d[I-C]) (Sigma), which helps reduce non-specific protein interactions with nucleic acids. The final volume of each footprint reaction was 11µl. First, a master mix of SaPI1 *stl-str* DNA was made as follows (per 10 total reactions); 10µl of 0.5pmol/µl SaPI1 *stl-str* DNA, 10µl of 1 µg/ µl poly(d[I-C]), 20µl of 5X EMSA buffer, and 2µl of 100mM CaCl2. 4 µl of SaPI1 *stl-str* DNA master mix was aliquoted into individual tubes on ice. 6µl of ∼3.3X Stl and/or Sri dilutions were made in EMSA buffer, pre-incubated on ice for 30 minutes, then added to 4 µl of SaPI1 *stl-str* DNA master mix, and allowed to incubate on ice for a further 30 minutes. To initiate DNase digestion, 1µl of DNaseI (2U/µl) was added to individual reactions and immediately transferred to a 30°C water bath for 10 minutes. Reactions were quenched by adding 17.2 µl of footprint loading buffer (0.5mL deionized formamide; 20µl of 0.25M EDTA, pH 7; 5µl XCFF and 5µl of BPB), followed by immediate transfer to dry ice. Samples were boiled at 100°C for 2 minutes immediately before loading onto a denaturing gel (5% polyacrylamide, 7M urea).

To help increase resolution of gel electrophoresis, denaturing gels were pre-electrophoresed in 0.5X TBE before adding samples (500V/400mA/400W for 30 minutes, then 750V/400mA/400W for 30 minutes; then 1,000V/400mA/400W for 30 minutes, and 1,250V/400mA/400W for > 30 minutes). 15µl of the quenched footprint reactions and the G/A ladder dilutions (discussed above) were loaded to gels. After samples were added to pre-electrophoresed gel, electrophoresis was continued for approximately 3,500 V-hrs for optimal resolution of the Stl-bound regions in SaPI1 *stl-str* DNA. G+A sequencing reactions were performed on the same end-labeled fragments using the method of Maxam and Gilbert (28) and run on the same gel to identify the sequences protected by Stl.

### Size-exclusion chromatography (SEC) and SEC small angle X-ray scattering (SEC-SAXS)

SEC analyses were carried out using a Superdex S200 26/600 column connected to an AKTA Pure system (GE Heatlhcare), equilibrated with buffer 100mM Tris-HCl pH 8, 300 mM NaCl and 5 mM MgCl2. Samples containing 200 μg of protein were loaded into the column and were eluted at a flow rate of 1 mg/ml.

SEC-SAXS was done on beamline B21 of the Diamond Light Source synchrotron (Didcot, UK). Data were recorded at 12.4 keV, at a sample-detector distance of 4.014 m using a Pilatus 2 M detector (Dectris, Switzerland). 50 µl of protein samples at concentrations of 10.0 mg/ml (Stl^SaPI1^) and 6.5 mg/ml (Stl^SaPI1^-Sri) were loaded onto a Superdex 200 Increase 3.2 size exclusion chromatography column in 10 mM Tris pH 8, 1% (w/v) sucrose at 0.075 ml/min using an Agilent 1200 HPLC system. The column outlet was fed into the experimental cell, and 620 x 3.0 s frames of SAXS data were recorded. Data were processed with ScÅtter IV (http://www.bioisis.net) as follows. The integral of ratio to background signal along with the estimated radius of gyration (Rg) for each frame was plotted. Frames within regions of low signal and low Rg recorded prior to protein elution were selected as buffer and subtracted from frames within regions of higher signal and constant Rg. Subsequent analysis was performed using the ATSAS 3.0 suite of programs (29). The radius of gyration Rg was obtained from the Guinier approximation following standard procedures. The pairwise distance distribution function p(r) was computed using the indirect Fourier transformation method implemented in GNOM (30). From the p(r) function, an alternative estimate of Rg and the maximum particle dimension Dmax were obtained. Molecular weights were estimated by Bayesian inference (31) in Primus. Ensemble optimization modelling was undertaken with EOM (22). 10000 models were generated in which the C-terminal domains (CTD, residues 101-247) were kept in the conformation observed by X-ray crystallography but the DBDs (residues 1-89) allowed to adopt positions consistent with their connection to the CTD via a native-like flexible linker (residues 90-100).

### Protein Crystallization and Data Collection

Crystals of 80*α* Sri-Stl^SaPI1^ SeMet derivative complex were obtained by vapor-diffusion technique using a sitting drop setup at 15°C in a reservoir solution of 0.4M Ammoniun phosphate, 25% PEG200. The crystals were cryo-protected using 30% of PEG200 solution when freezing in liquid nitrogen. Single-wavelength anomalous diffraction (SAD) on the I03 beamline at the Diamond Light Source synchrotron radiation facility (DLS; Didcot, UK) (32) was collected at a wavelength of 0.98 and at 2.8 Å. Data were indexed, integrated, and scaled using the program autoPROC (33). Model building and refinement was performed with CCP4 suite and Coot (34, 35). The crystallographic parameters, data-collection and refinement statistics are listed in Table 1.

### Characterization of the SaPI1 *str* promoter

To characterise the *str* promotor, RNA extraction using Ambion kit (Novartis) was performed using the RN4220 strain lysogenic for 80⍺ and carrying SaPI1, after 90 min of phage induction with 2 μg/ml MC. 5’/3’RACE Kit (Roche) was used to amplify the RNA obtained and the final DNA was sequenced to obtain the +1 nucleotide of *str.* The -10 and -35 RNA pol binding sites were localised after the analysis of the DNA sequence.

### Electrophoretic Mobility Shift Experiments (EMSA)

The Stl DNA binding regions for the EMSA experiments were obtained commercially and hybridized at 95°C 15 min with equal concentration of the forward and reverse primers (Table S8). Final DNA concentration of 1μM and increasing concentrations of Stl (0.5-8 μM) were mixed with 1μg/ml poly(d[I-C]) (Roche) in EMSA buffer (50 mM Tris-HCl pH 8, 5 mM MgCl2, 1mM DTT, 0.1 mM EDTA, and 5% glycerol) and incubated for 30 min at room temperature.

The samples were then load in 6% Tris-Borate-EDTA (TBE) polyacrylamide gels were electrophoresed in TBE buffer at 90 V for 1-2 h. Gels were stained with Gel Red (Biotium) for 10min in shaking conditions and analysed by ChemiDoc (BioRad).

### Pull-down experiments

The wt and the Y76A versions of the Stl^SaPI1^-Sri complex, cloned in the pPROEX-Hta plasmid, were expressed in 20 ml of culture as previously indicated. The cells where then lysed with BugBuster protein extraction reagent (Novagen) for 30 min at room temperature. The soluble fraction was then incubated 1h with HisPur Ni-NTA Resin (Thermofisher), and the proteins bound to the resin purified as described before. After purification the fractions were analyzed by SDS-PAGE, stained with Instant Blue (Expedeon) and visualized with ChemiDoc visualizer (BioRad).

### β-Lactamase assays

Cells were obtained at different time points after MC induction of the lysogenic cells carrying the appropriate plasmids. β-lactamase assays, using nitrocefin as substrate, were performed as described (5, 4). Briefly, 50 µl of the collected sample were mixed with 50 µl of nitrocefin stock solution (192 µM made in 50 mM potassium phosphate buffer, pH 5.9), and immediately reading the absorbance at 490 nm using a FLUOstar Omega (BMG LABTECH) for 45 min. Promoter activity was calculated as Promoter activity = (*d*A490/*d*t(h))/(OD540 x d x V), where OD540 is the absorbance of the sample at OD540 at collection, d is the dilution factor, and V is the sample volume.

### SaPI transfer

*S. aureus* strains lysogenic for helper phages and containing the required SaPI(s) were grown to early exponential phase (OD540∼0.15) at 37°C and 120 rpm. Cultures were then induced by the addition of mitomycin C (2 µg ml^-1^) and incubated for 4-5 h at 30°C followed by overnight incubation at room temperature before filtering the lysate with a 0.2 µm syringe filter. For SaPI titre determination, *S. aureus* RN4220 strain was grown overnight at 37°C and 120 rpm. The culture OD was adjusted to OD540∼1.4 with TSB and supplemented with 4.4 mM CaCl2. 100 µl of the appropriate lysate dilution were added to 1 ml of this cell suspension and incubated for 20 min at 37°C. Three ml of transduction top agar (TTA, 30 g l^-1^ TSB, 7.5 g l^-1^ agar) were added to the transduction and the mix poured onto a TSA plate containing the appropriate antibiotic. Plates were incubated for 16-24 at 37°C prior to determination of transducing units.

### Quantification and statistical analysis

All statistical analyses were performed as indicated in the figure legends using GraphPad Prism 6.01 software.

### Data Availability

Atomic coordinates and structure factors have been deposited at the RCSB Protein Data Bank (PDB code 7P4A).

## ACKNOWLEDGEMENTS

We thank Kristin Lane and Jamie Brooks for creating the expression plasmids pKDL97 and pJLB20, respectively. We also thank Deborah Hinton for her assistance with the footprinting experiments. This work was supported by grants MR/V000772/1, MR/M003876/1 and MR/S00940X/1 from the Medical Research Council (UK), BB/N002873/1, BB/S003835/1 and BB/V002376/1 from the Biotechnology and Biological Sciences Research Council (BBSRC, UK), Wellcome Trust 201531/Z/16/Z, and ERC-ADG-2014 Proposal n° 670932 Dut-signal from EU to J.R.P.; grants PID2019-108541GB-I00 from Spanish Government (Ministerio de Economía y Competitividad y Ministerio de Ciencia e Innovación) and PROMETEO/2020/012 from Valencian Government to A.M.; grants MOE2017-T2-2-163 and MOE2019-T2-2-162 from the Ministry of Education to J.C.; and grant NIH R01 AI083255 to G.C. J.T. was supported by NIH IRACDA Grant K12GM093857 to Virginia Commonwealth University. . L.M-R was the recipient of a Spanish postdoctoral fellowship from Fundación Ramón Areces (2018-2020). J.R.P. is thankful to the Royal Society and the Wolfson Foundation for providing him support through a Royal Society Wolfson Fellowship.

## AUTHOR CONTRIBUTIONS

A.M. and J.R.P. conceived the study. L.M-R., M.A., J.B., J.T., and O.B. conducted the experiments. L.M-R., M.A., J.B., J.T., R.J.C., J.C., O.B., G.E.C, A.M. and J.R.P analysed the data. G.E.C and J.R.P wrote the manuscript.

## DECLARATION OF INTERESTS

The authors declare no competing interests.

---

**Table S1.**
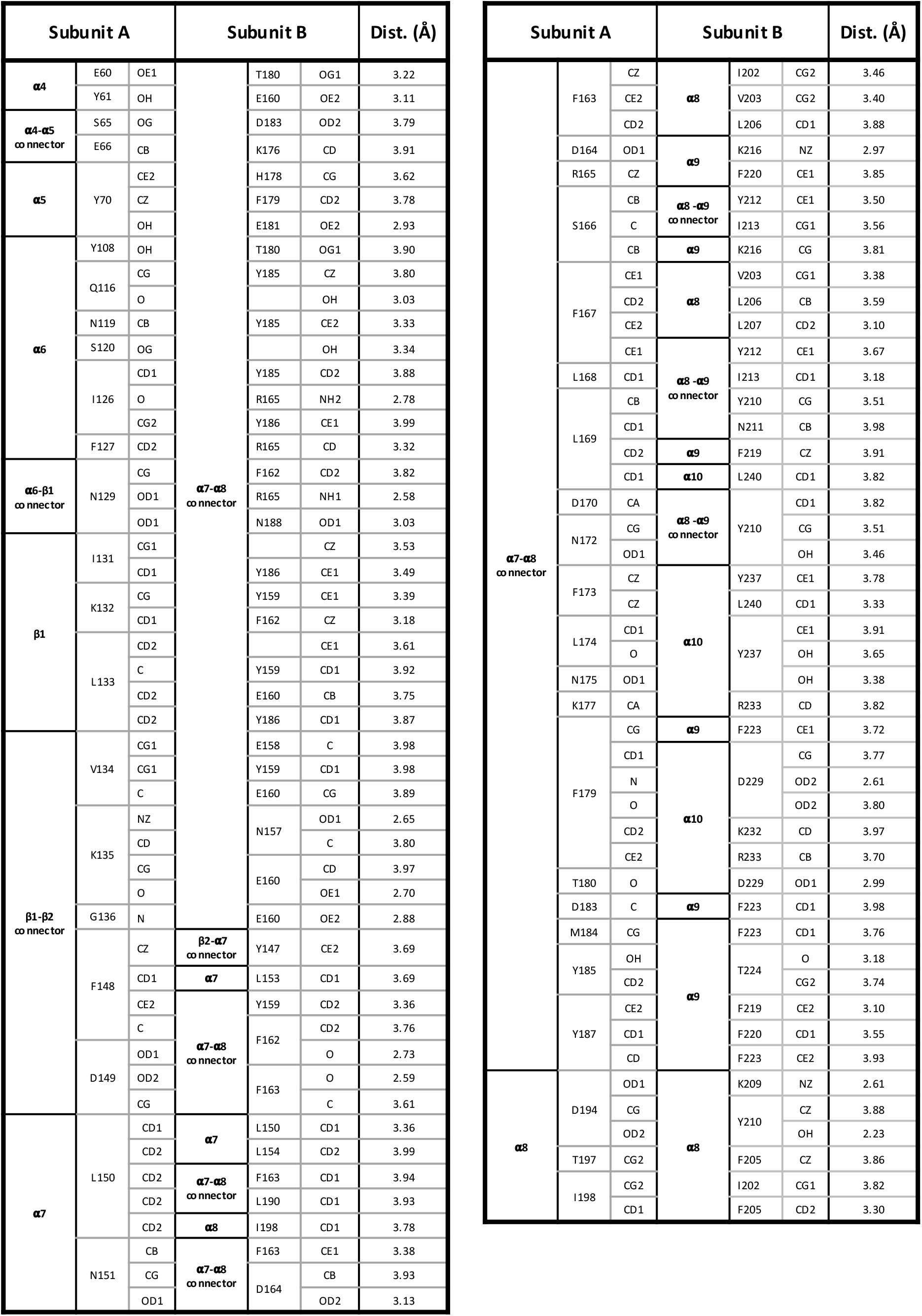
Stl^SaPI1^ dimer contacts.

**Table S2.**
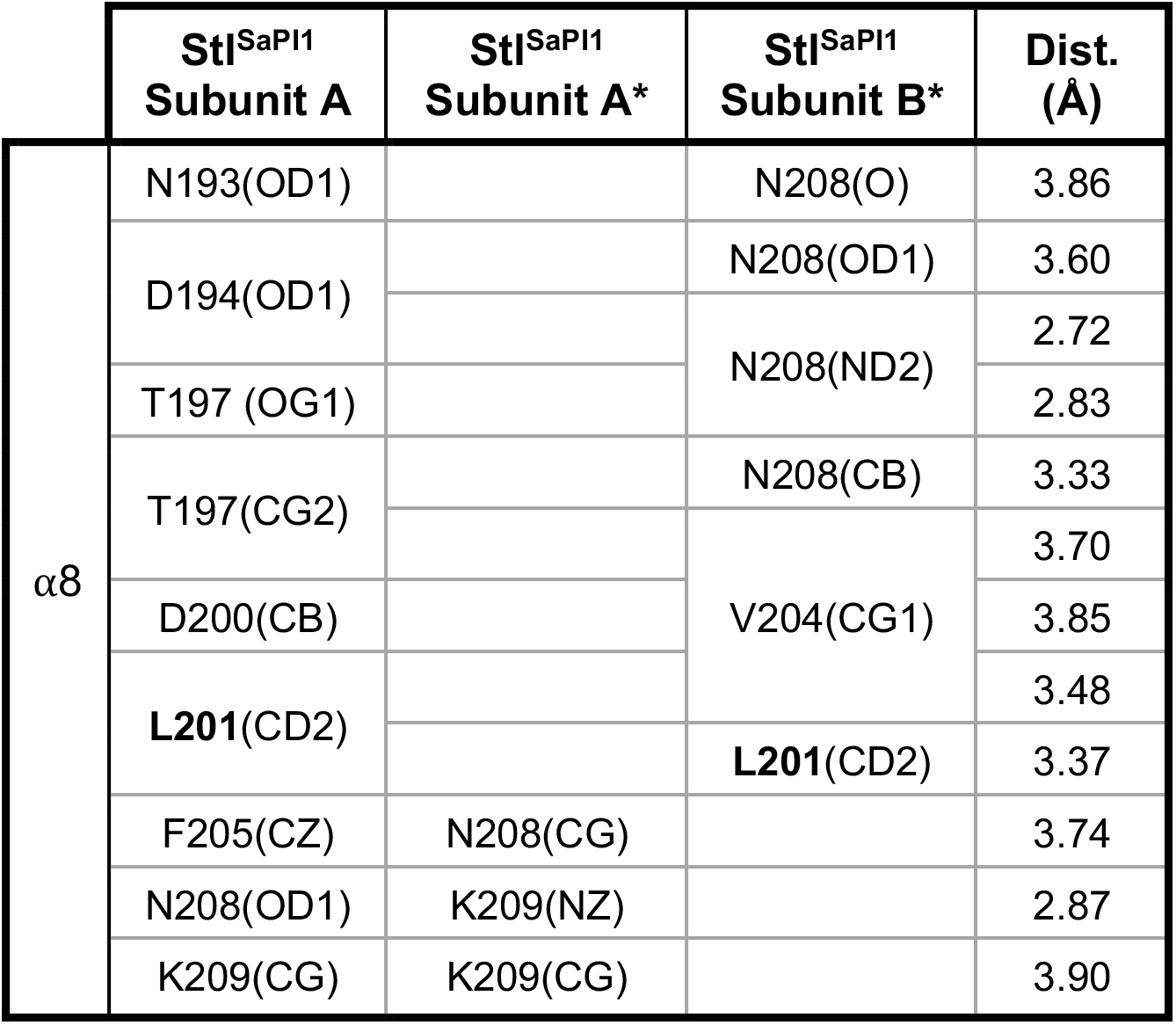
Stl^SaPI1^ tetramer contacts.

**Table S3.**
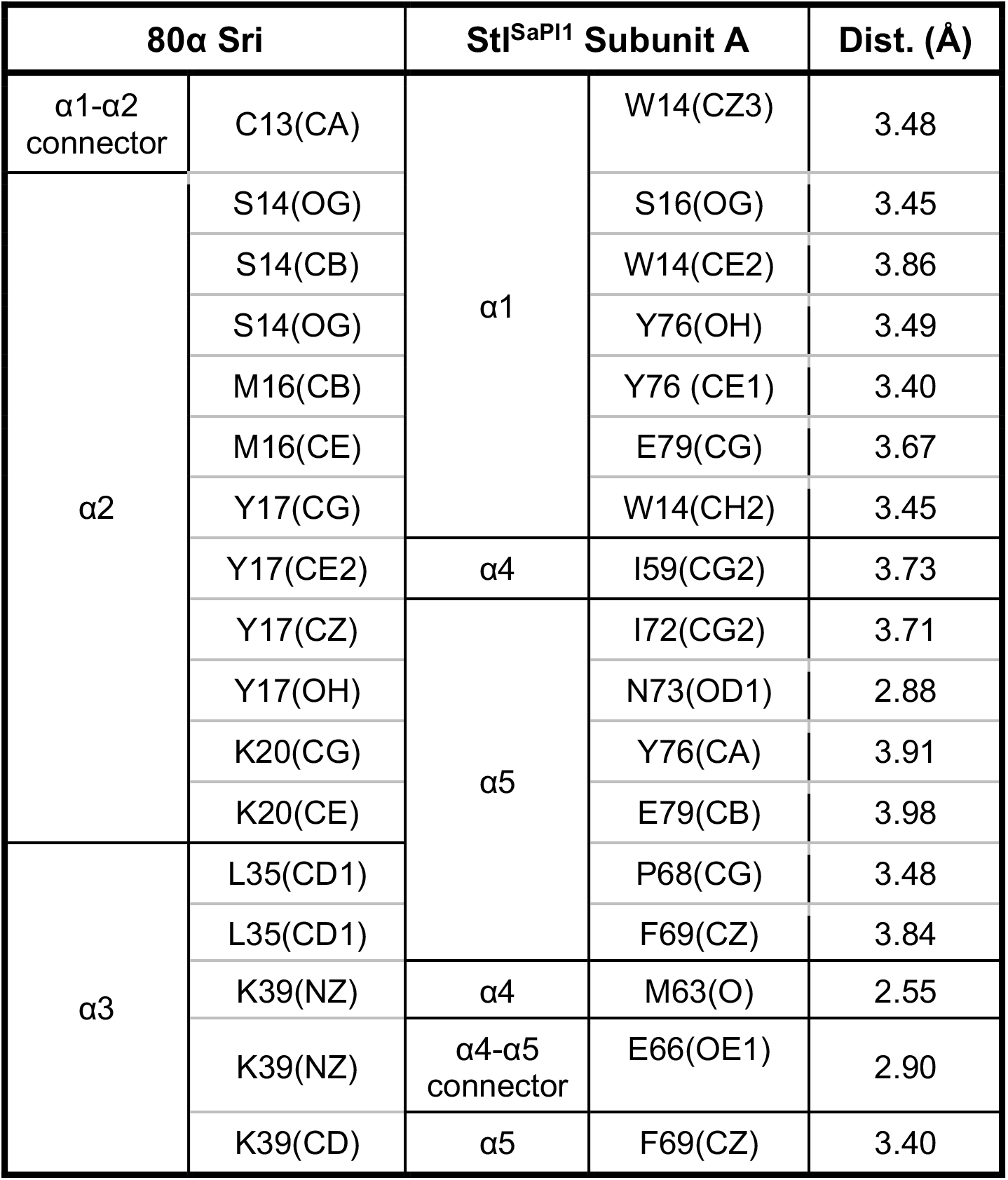
Stl^SaPI1^ –Sri contacts.

**Table S4.** Stl^SaPI1^-like repressors.

| Name | Strain | Protein accession number | Gen accession number (Genomic location) |
| --- | --- | --- | --- |
| Stl<sup>SaPI1</sup>-like SaPI | *Staphylococcus aureus* NTCT9546 | WP_115208074.1 | NZ_UHAK01000002.1 (417268-417987) |
| Stl<sup>SaPI1</sup>-like ShPI | *Staphylococcus hominis* FDAARGOS | SIH39397.1 | FSLI01000015.1 (137382-137948) |
| Stl<sup>SaPI1</sup>-like V.pan | *Virgibacillus panthothenticus* | WP_077296842.1 | NZ_Lt727679.1 (439288-439887) |
| Stl<sup>SaPI1</sup>-like B.sp | *Bacillus* sp. X1 | WP_144546869.1 | NZ_VECT01000031.1 (61128-61724) |
| Stl<sup>SaPI1</sup>-like V.sp | *Virgibacillus* sp. SK37 DE0373 | WP_145524117.1 | NZ_VDZE01000056.1 (8525-9187) |
| Stl<sup>SaPI1</sup>-like B.cana | *Bacillus canaveralius* ATCC29669 | WP_101577990.1 | NZ_PGVA01000029.1 (45243-45830) |
| Stl<sup>SaPI1</sup>-like B.encl | *Bacillus enclensis* CARE-V7 29 | MBH9968881.1 | NZ_JADZLX100000029.1 (12298-12888) |

**Table S5.** Bacterial strains used in this work.

| Stock | Description | Reference |
| --- | --- | --- |
| DH5α | *E. coli* host for DNA cloning | Invitrogen |
| RN4220 | *S. aureus* restriction-defective derivate of RN450 | Lab strain |
| BL21 (DE3) | *E. coli* expression strain | Stratagene |
| JP19607 | *E. coli* DH5α pJP2371 | This study |
| JP22989 | *E. coli* DH5α pJP2700 | This study |
| JP709 | *E. coli* DH5α pJP2344 | Lab strain |
| JP17789 | *E. coli* DH5α pJP2366 | This study |
| JP17792 | *E. coli* DH5α pJP2369 | This study |
| JP17813 | *E. coli* DH5α pJP2387 | This study |
| JP17816 | *E. coli* DH5α pJP2390 | This study |
| JP2799 | RN10359 lysogenic for wt 80α | Lab strain |
| JP17848 | RN4220 Y76A StI SaPI1 | This study |
| JP19902 | RN10359 lysogenic for wt 80α + Y76A StI SaPI1 | This study |
| JP17863 | RN4220 pJP2387 | This study |
| JP17866 | RN4220 pJP2390 | This study |
| JP17891 | JP2799 pJP2387 | This study |
| JP17894 | JP2799 pJP2390 | This study |
| JP15526 | BL21 (DE3) pJP2344 | This study |
| JP17839 | BL21 (DE3) pJP2366 | This study |
| JP17842 | BL21 (DE3) pJP2369 | This study |
| JP17873 | RN4220 pCN41 | This study |
| JP17901 | JP2799 pCN41 | This study |
| JP22095 | *E. coli* DH5α pJP2699 | This study |
| JP22101 | JP2799 pJP2699 | This study |

**Table S6.** Plasmids used in this work.

| Plasmid | Description | Reference |
| --- | --- | --- |
| pPROEX-Hta | Expression vector | Invitrogen |
| pCN41 | Expression vector | (36) |
| pJP2700 | pPROEX Hta His-Stl SaPI1 | This study |
| pJP2344 | pPROEX Hta His-Sri 80α + Stl SaPI1 | This study |
| pJP2366 | pPROEX Hta His-Sri 80α + Y76A Stl SaPI1 | This study |
| pJP2369 | pPROEX Hta His-Sri 80α + L201E Stl SaPI1 | This study |
| pJP2371 | pPROEX Hta His-L201E Stl SaPI1 | This study |
| pJP2384 | pCN41- Stl-Str region of SaPI1 | This study |
| pJP2387 | pCN41- (Y76A Stl)-Str region of SaPI1 | This study |
| pJP2390 | pCN41- (L201E Stl)-Str region of SaPI1 | This study |
| pJP2699 | pCN41- Stl-Str region of SaPI1 with *Pstr* -35 mutant (TTGACA to TTAACA) | This study |
| pKDL97 | pGEX-4T-1 (GE Healthcare) derivative with 80□ *sri* inserted between the *Bam*HI and *Xho*I sites to generate an N-terminal GST fusion. | This study |
| pJLB20 | pET15b (Novagen) carrying the SaPI1 *stl* gene inserted between the *Nde*I and *Bam*HI sites to generate an N-terminal His tag. | This study |

**Table S7.**
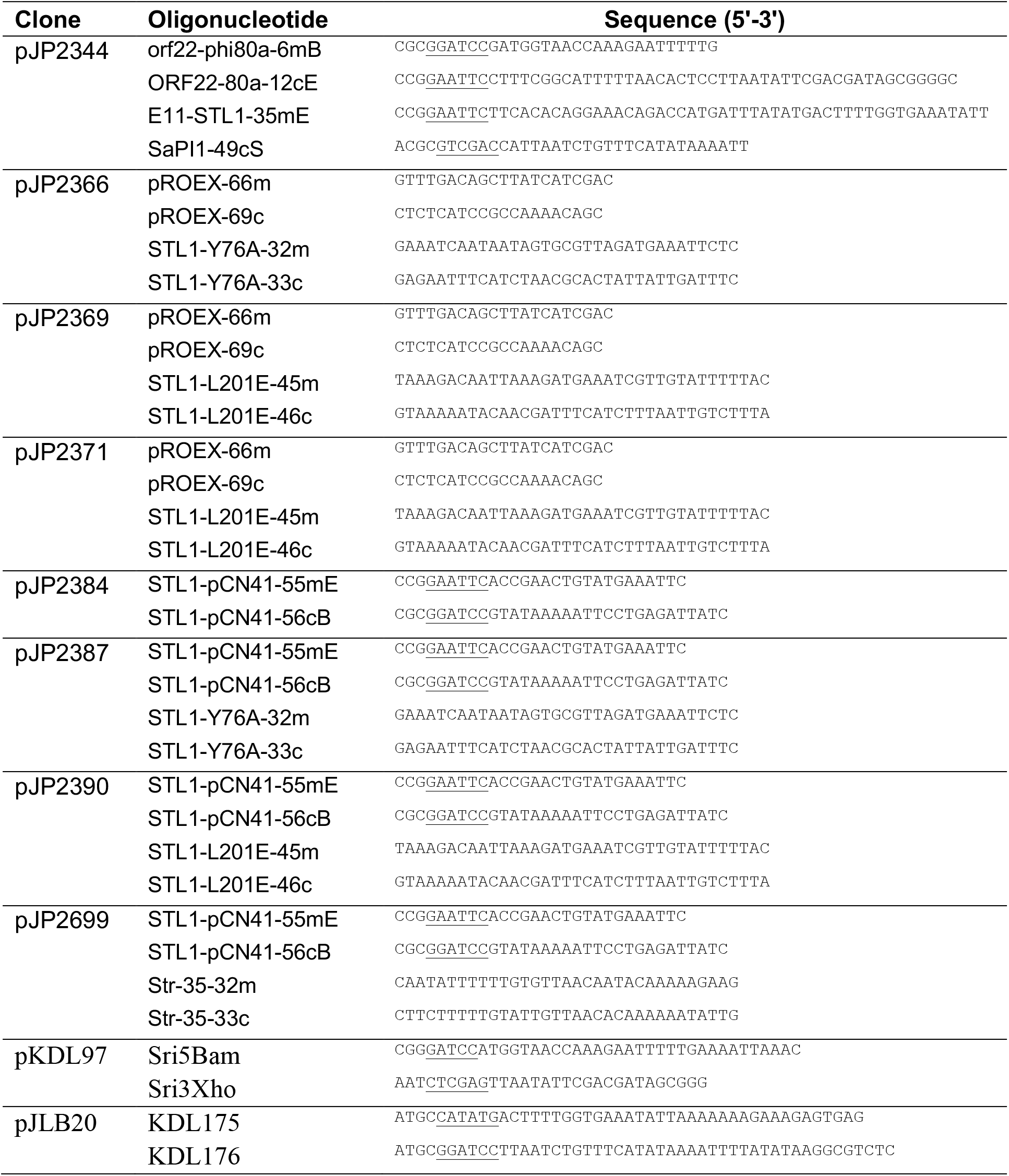
Primers used in this work.

**Table S8.**
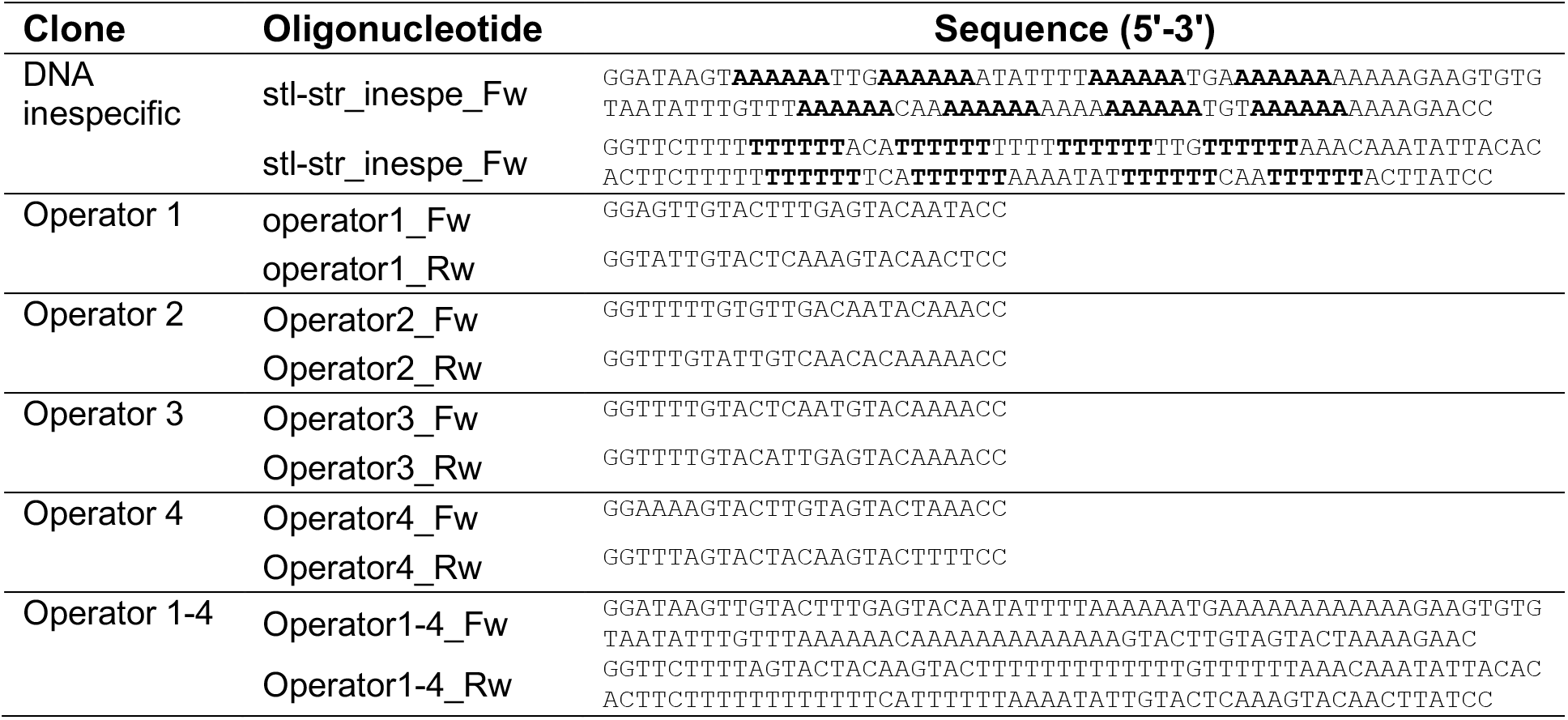
Primers used for EMSA in this work.

**Figure S1.**
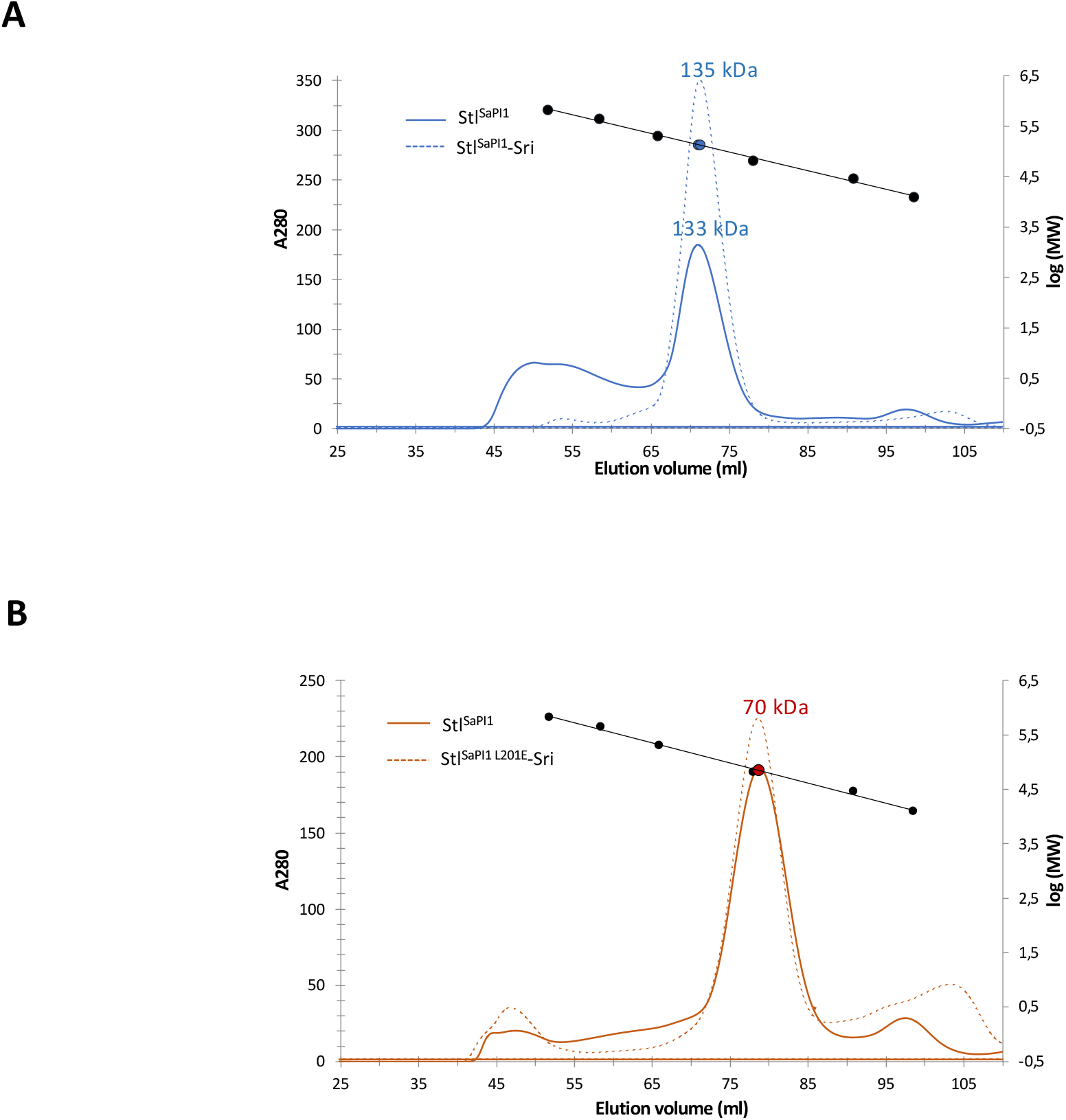
Stl^SaPI1^ oligomeric state. Size exclusion experiments with wt Stl^SaPI1^ (A) and Stl^SaPI1 L201E^ (B), both alone (solid lines) or in complex with Sri (dashed lines). The molecular weight for the samples (blue and red dots) and for the calibration proteins (black dots) are represented in the right *y* axes in both graphs.

**Figure S2.**
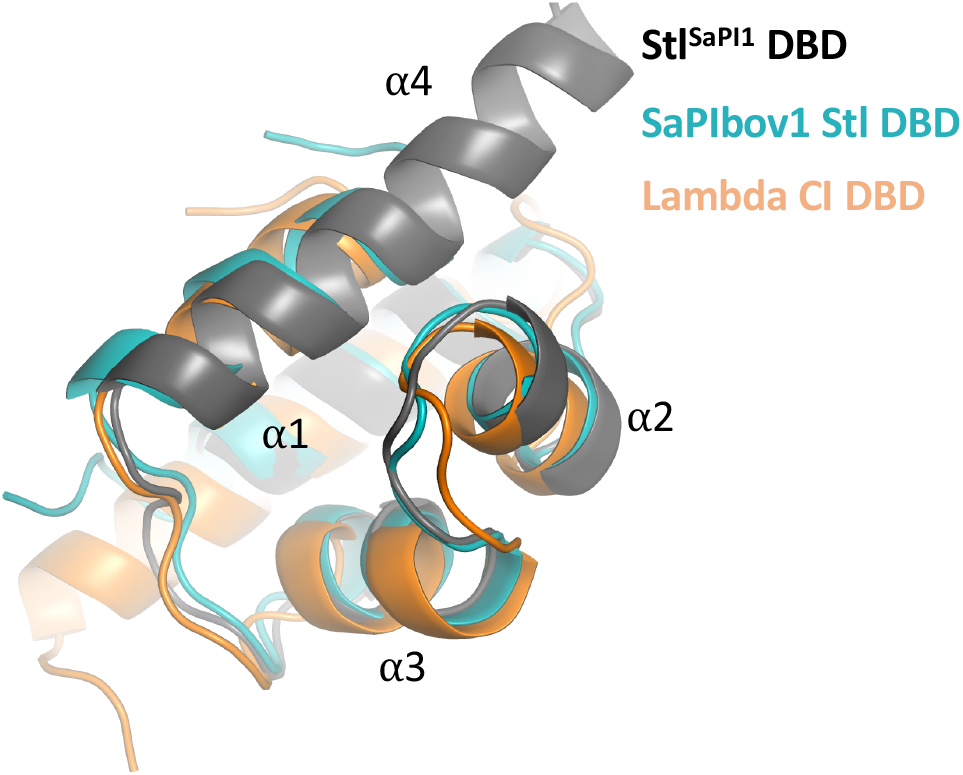
HTH family DBD comparison. Superimposition of Stl^SaPI1^ DBD in black, SaPIbov1 Stl DBD (PDB 6H49) in blue, and CI lambda phage repressor DBD (PDB 1LMB) in orange. The four alpha helices are marked.

**Figure S3.**
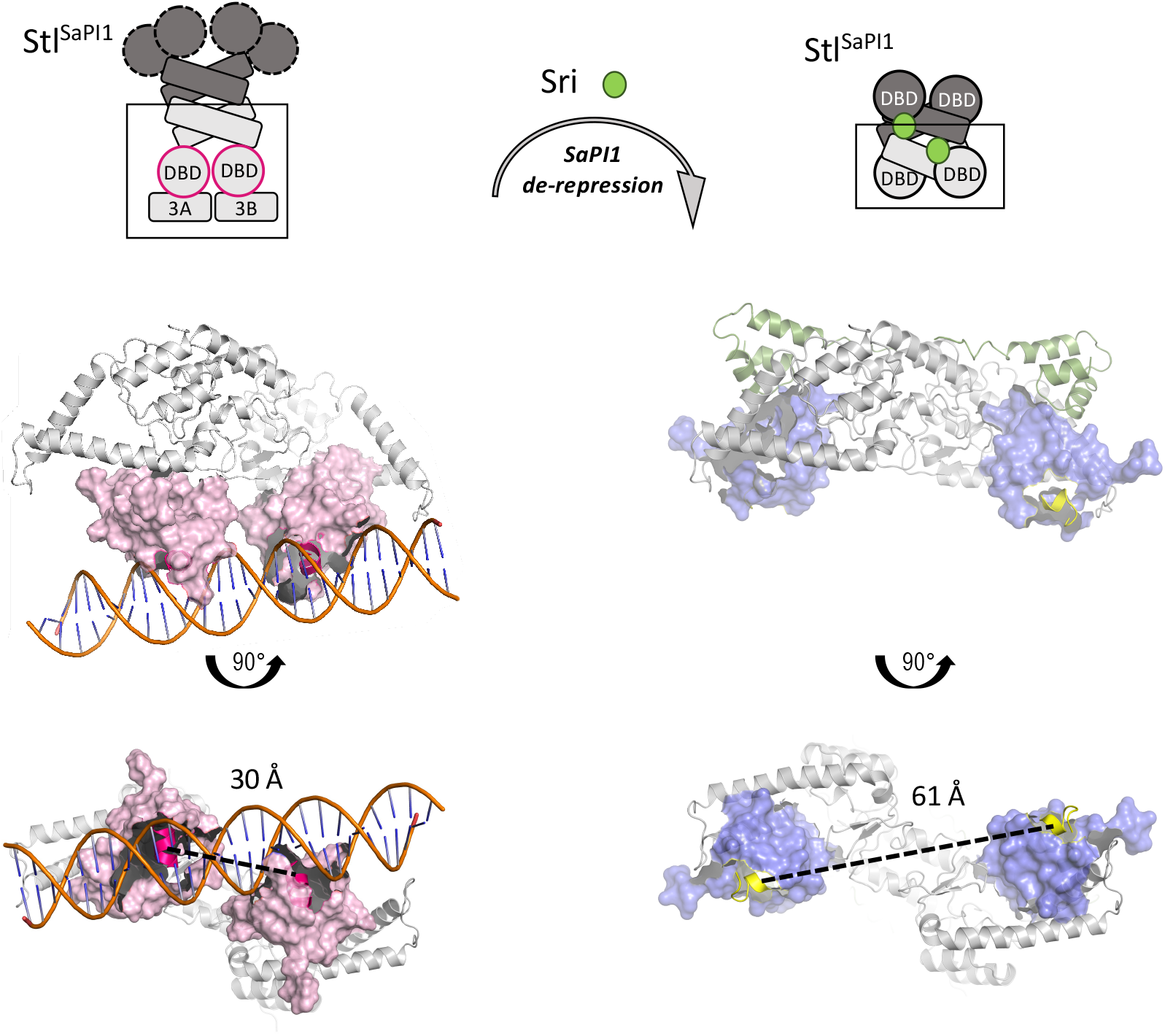
Stl^SaPI1^ DBD flexibility. Models for the location of the Stl^SaPI1^ DBDs when the repressor is bound to its operator regions repressing SaPI1 (left), or when Stl^SaPI1^ is bound to Sri, derepressing SaPI1 (right). The DBDs are represented as surface in pink for Stl^SaPI1^-DNA and in blue for Stl^SaPI1^-Sri complexes. Sri is represented in green. The DBD ⍺3 helix responsible for the interaction with the DNA is represented in cartoon in dark pink for Stl^SaPI1^-DNA and in yellow for Stl^SaPI1^-Sri. The distance between ⍺3 helices in the Stl^SaPI1^ dimer is indicated in the figure.

**Figure S4.**
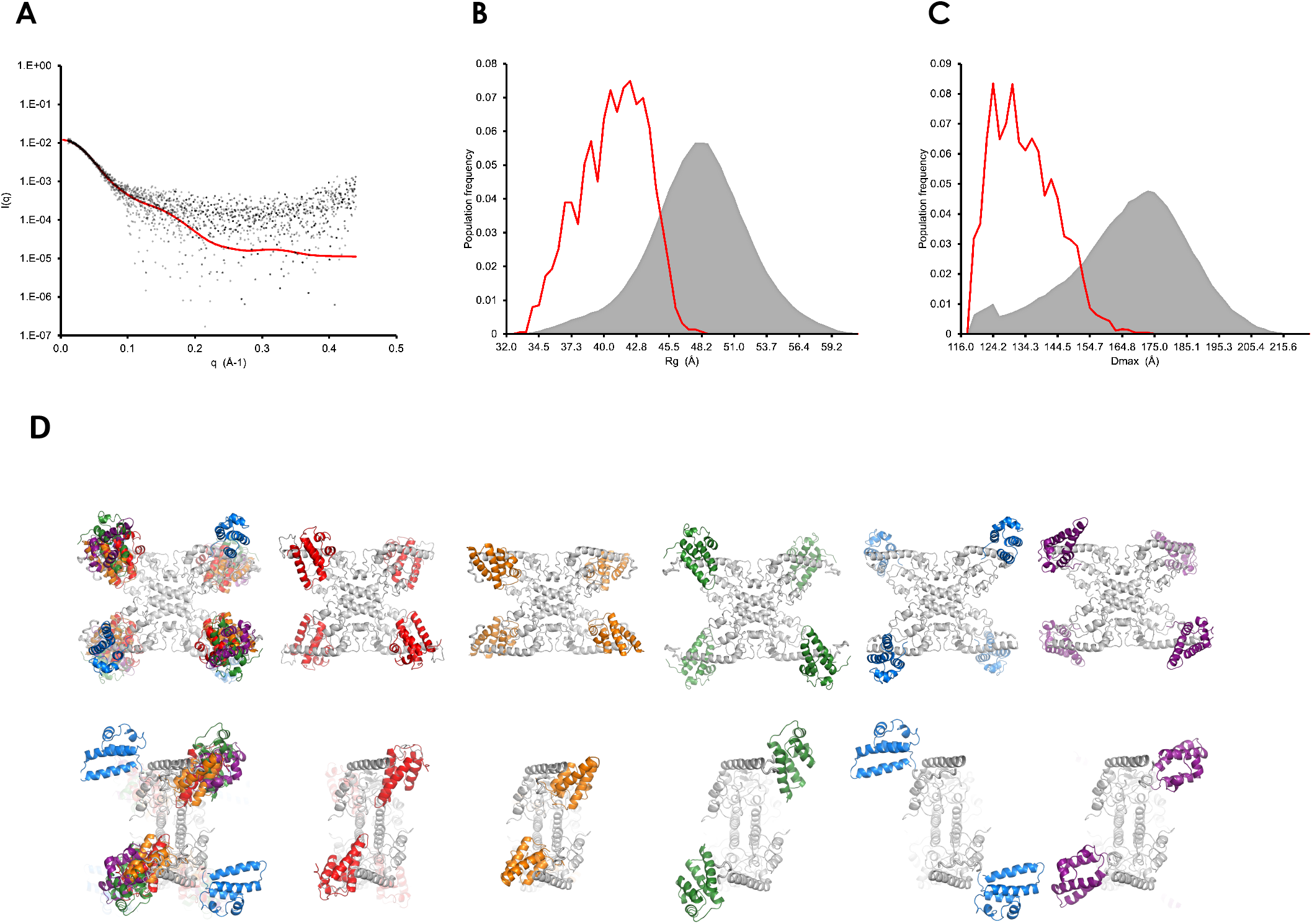
SAXS analysis of Stl^SaPI1^ tetramer conformational heterogeneity. (**A**) Overlay of experimental SAXS data for Stl^SaPI1^ (dots) and those computed for the best-fitting ensemble selected by EOM with χ^2^ = 1.68 (red line). (**B**) Rg and Dmax (C) computed for a pool of 10000 solution conformers (grey) compared with that computed for the best-fitting ensemble generated by the ensemble optimization method (EOM) in red for Stl ^SaPI1^ tetramer in which the C-terminal domains were kept in the conformation observed by X-ray crystallography but the N-terminal DNA-binding domains were allowed to adopt positions consistent with their connection to the CTD via a native-like flexible linker. (**D**) Orthogonal views of the superimposition upon a model for the Stl^SaPI1^ tetramer of 4 Stl^SaPI1^ conformers that combined provide the best fit to the SAXS data via the EOM. The fit was obtained with ∼ 50% of model 1 (DBDs in orange), 25% of model 2 (DBDs in green), and 13% of models 3 (DBDs in blue) and 4 (DBDs in purple). DBDs of the Stl^SaPI1^ tetramer model are in red.

## Notes

### Competing Interest Statement

The authors have declared no competing interest.

